# Spatial Partitioning of Glucose Metabolism Supports Tissue-Specific Metabolic Programs In Vivo

**DOI:** 10.1101/2025.11.09.687253

**Authors:** Ian J. Gonzalez, Aaron D. Wolfe, Benjamin Clark, Michael Hanna, Gabrielle E. Geise, Quishi Sun, Snusha Ravikumar, Anastasia Tsives, Sarah E. Emerson, Stephan Siebel, Richard Kibbey, L. Safak Yilmaz, Albertha J.M. Walhout, Daniel Colón-Ramos

## Abstract

Tissues exhibit metabolic heterogeneity that tailors conserved pathways to distinct physiological demands, yet how this heterogeneity is achieved *in vivo* remains poorly understood. Here, we use *Caenorhabditis elegans* to investigate tissue-specific requirements for glucose-6-phosphate isomerase (GPI-1), a conserved reversible enzyme that links glycolysis and the pentose phosphate pathway (PPP). Tissue-specific metabolic-network modeling predicted differential glycolytic and PPP flux potential across adult tissues and identified tissue-specific biases in GPI-1 reaction directionality. Genetic disruption of *gpi-1* produced germline defects consistent with impaired PPP-associated anabolic metabolism and somatic defects consistent with impaired glycolysis, indicating that GPI-1 supports distinct metabolic functions across tissues. We further discovered that two GPI-1 isoforms are differentially expressed and localized: GPI-1A is broadly expressed and cytosolic, whereas GPI-1B is enriched in the germline and localizes to endoplasmic reticulum–associated compartments. Isoform-specific perturbations revealed distinct requirements for GPI-1A and GPI-1B in somatic glycolysis and reproductive physiology. These findings implicate isoform-specific subcellular localization as a possible contributor to the partition of the functions of a conserved reversible enzyme, enabling tissue-specific anabolic and catabolic metabolism *in vivo*.

## INTRODUCTION

The processing of glucose molecules in the early reactions of glycolysis determines whether intracellular glucose generates energy or is diverted into biosynthetic pathways such as the pentose phosphate pathway (PPP)^1–3^. Glucose metabolism therefore sits at the crossroads of anabolic and catabolic demands that are shared across cells yet differentially tuned to the needs of individual cell types. Consistent with this, the enzymes that regulate upper glycolysis are broadly conserved and present across tissues, yet their relative contributions to glucose utilization differ across organisms, cell types, and even subcellular compartments, shaping how carbon is allocated toward anabolic or catabolic pathways in specific contexts^4–6^. This differential regulation has physiological significance, as illustrated by tissue-specific differences in glucose flux through glycolysis versus the PPP in metazoans. For instance, biochemical approaches using isotope-labeled glucose demonstrated that in neurons and hepatocytes, glucose 6-phosphate (G6P), the product of the first step in upper glycolysis, typically proceeds toward lower glycolysis rather than the PPP, with reported flux ratios as high as 40:1^7–9^. In contrast, in adipose and ovarian tissues, these ratios can drop to ∼2:1, reflecting a shift toward PPP activity and anabolic carbon use^10–12^. This metabolic heterogeneity mirrors distinct physiological demands: Neurons rely on rapid glycolytic ATP production to fuel transient bursts of activity^13–17^, whereas lipogenic tissues require NADPH that can be produced by the PPP to sustain fatty acid synthesis^18–20^. While these studies demonstrate tissue-specific metabolic heterogeneity in glucose utilization in animals, how these tissue-specific patterns of glucose metabolism are established *in vivo* is poorly understood.

Differential metabolism across tissues could arise from the expression of tissue-specific isoforms of metabolic enzymes. Consistent with this hypothesis, transcriptomic and proteomic studies have revealed glycolytic enzyme isoform expression that is specific to certain tissues^21–24^. Biochemical studies have also demonstrated that different glycolytic enzyme isoforms, that share high degrees of sequence similarity, display distinct modes of regulation. For example, phosphofructokinase M (PFKM) is the predominant isoform in skeletal muscle and is expressed at low levels in other tissues. *In vitro* reconstitution and activity assays show that, relative to the other PFK isoforms (PFKL and PFKP), PFKM displays higher enzymatic activity and reduced inhibition by ATP, rendering it less regulated but more efficient at sustaining glycolytic flux^22,25–30^. The physiological importance of such tissue-specific enzyme expression is evident in human disorders: mutations specifically in *PFKM* cause glycogen storage disease, muscle dysfunction, and erythrocyte hemolysis^31–33^. A gap in knowledge remains, however, in how distinct isoforms of metabolic enzymes are distributed *in vivo* and how their distribution might contribute to tissue-specific functions in an intact system.

This regulation could be particularly important at metabolic pathway branch points that direct glucose utilization towards anabolism or catabolism. The glycolytic enzyme glucose-6-phosphate isomerase (GPI) catalyzes the reversible interconversion between G6P and fructose-6-phosphate (F6P), a reaction that biochemically connects the entry points of glycolysis and the PPP. This reversible interconversion of G6P to F6P is near equilibrium, and in biochemical assays GPI allows flux in either direction depending on the relative concentrations of G6P and F6P^5^. Yet in specific tissues, flux through GPI is regulated towards specific directions. For example, immune cells display the capacity to shift flux toward anabolic reactions through the PPP to generate reducing power for oxidative bursts during immune responses, whereas neurons bias flux toward catabolic reactions through glycolysis to sustain synaptic transmission during periods of activity^34,35^. The non-oxidative arm of the PPP can also regenerate F6P from pentose phosphates, enabling a cyclic mode of PPP activity that may depend on reverse GPI flux to recycle F6P back to G6P^34^. Whether GPI flux is regulated in a tissue-specific manner *in vivo* to bias reaction directionality remains unknown.

Addressing how branch-point enzymes such as GPI regulate glucose flux between glycolysis and PPP *in vivo* has been challenging, as genetic ablation or molecular manipulation of these essential metabolic genes often leads to lethality. Moreover, many glycolytic and PPP enzymes are encoded by redundant isoforms in metazoans, complicating efforts to dissect their individual contributions to tissue-specific metabolism. To overcome these limitations, we established a system in *Caenorhabditis elegans* that is uniquely suited to examine these relationships under physiological conditions. *C. elegans* offers several key advantages: (1) conservation of metabolic genes and pathways^36^; (2) reduced redundancy in the number of genes encoding proteins regulating specific steps^37^; (3) metabolic network models (MNM) of tissue-specific metabolism20,38–40, (4) genetic mutants to precisely manipulate the enzymatic steps *in vivo*^15^; (5) quantifiable physiological readouts of the emerging phenotypes in intact, living animals; and (6) tissue-specific metabolic sensors to spatially resolve and estimate flux^15,41^. By integrating genetic engineering, metabolic-network modeling, metabolic profiling, quantitative physiology, and cell biological analysis, this system enables dissection of the differential roles, regulatory modes, and vulnerabilities of glycolysis and the PPP at single-tissue resolution *in vivo*.

In this study we use *C. elegans* to dissect how a single reversible glycolytic reaction supports divergent metabolic demands across tissues. We find that tissue-specific physiological roles are supported by divergent mechanisms of GPI-1 function: as a PPP enzyme in the biosynthetically-demanding germline, and as a classical glycolysis enzyme in somatic tissues during energetically-demanding contexts. Additionally, although GPI-1 is broadly expressed *in vivo*, our findings reveal tissue-specific differences in GPI-1 isoform expression and suggest that distinct isoforms might support different aspects of GPI-1 function. Together, these findings suggest that tissue-specific isoform expression and subcellular localization can diversify the functions of conserved reversible metabolic enzymes to support distinct anabolic and catabolic programs in specific tissues *in vivo*.

## RESULTS

### Metabolic network modeling predicts tissue-specific routes of glucose metabolism

To gain insight into how different tissues may preferentially execute glycolysis or the pentose phosphate pathway (PPP), we extended our recent whole-animal metabolic wiring analysis of *C. elegans*^20^ to specific tissues. In that study, we identified a predicted cyclic PPP flux mode, but the whole-animal model did not specify which tissues contributed to this metabolic program. To localize this predicted flux pattern and identify potential tissue-specific roles, we analyzed single-cell RNA-seq data from Day 1 adult animals across 18 cell types^38,42^. Specifically, we integrated these transcriptomic data with iCEL1314, the genome-scale metabolic network model of *C. elegans*, using the MERGE pipeline to predict reaction flux carrying capacities and reaction directionality for each cell type38,42 (Fig 1A).

**Figure 1:**
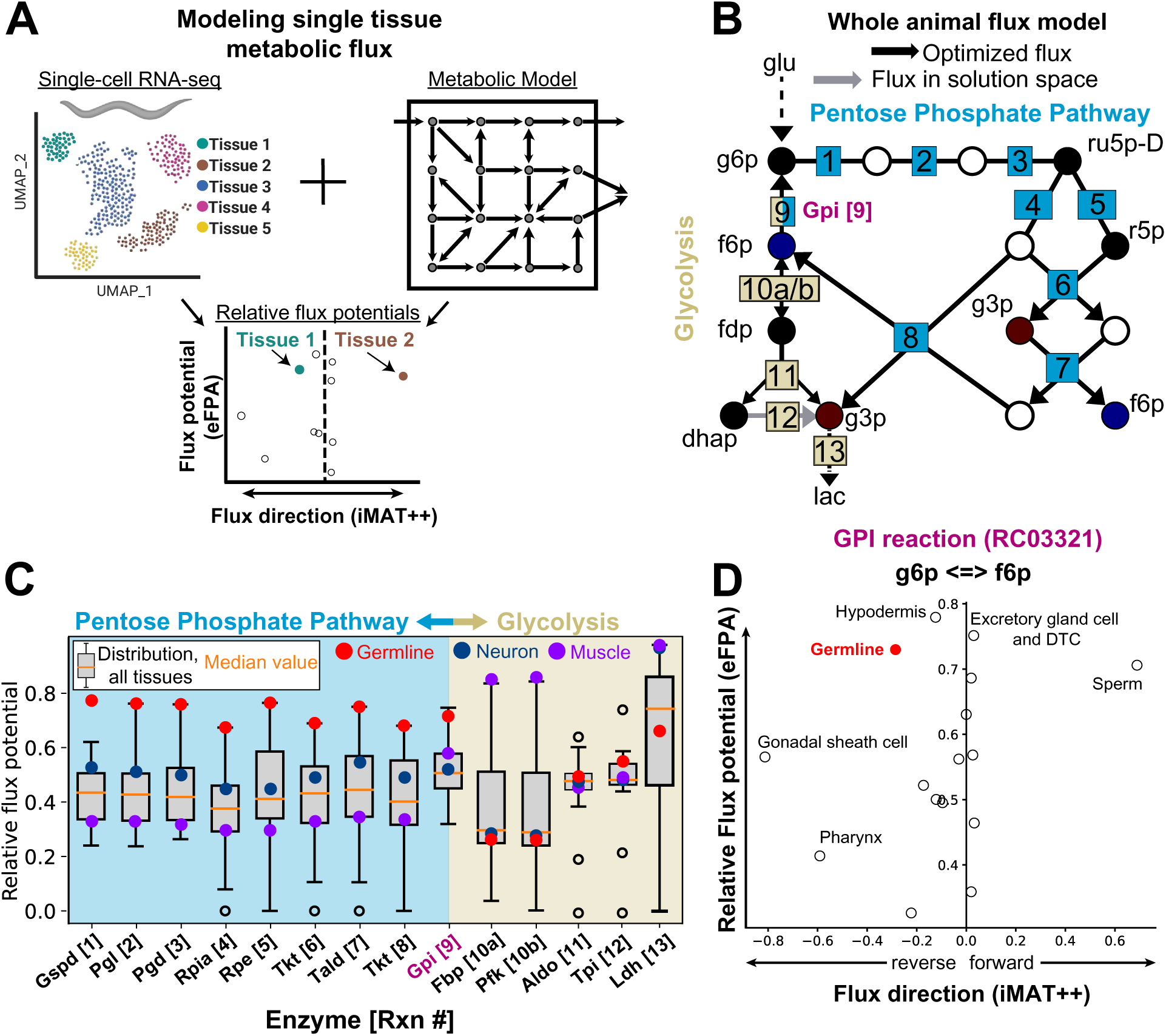
Metabolic network modeling predicts tissue-specific patterns of carbohydrate metabolism. **(A)** Pipeline diagram for integration of single-cell RNA-seq data from Day 1 adult worms^42^ with a metabolic network model that enables tissue-specific flux predictions for individual metabolic reactions. **(B)** Whole animal flux model from metabolic wiring analysis of *C. elegans*^20^ predicting reverse flux through the GPI-1 reaction (denoted in purple as reaction 9) followed by flux of G6P into the PPP. (C) Relative flux potentials for the reactions in (B), with glycolysis and PPP reactions colored gold and blue, respectively. Box plots represent distribution of relative flux potential values across all tissues. Reaction numbers correspond to those shown in (B). **(D)** Relative flux potentials predicted by the eFPA algorithm plotted against fluxes predicted by the iMAT++ algorithm for the GPI-1 reaction (RC03321). The x-axis indicates predicted flux state: absence/presence (zero vs. nonzero) and direction (sign), where negative values denote reverse flux (F6P → G6P; left) and positive values denote forward flux (G6P → F6P; right).

We identified tissue-level differences in the predicted relative flux-carrying capacity of reactions in the PPP and upper glycolysis (Fig. 1B). These predictions were based on relative flux potential (rFP) scores generated by enhanced Flux Potential Analysis (eFPA), which integrates local gene expression levels for each reaction with expression information from surrounding pathways^39^ (Table S1). rFP predictions for PPP and upper glycolytic reactions varied significantly across tissues (Fig. 1C). For example, the germline showed high predicted rFP across PPP reactions, whereas somatic tissues such as muscle showed high predicted rFP across glycolytic reactions (Fig. 1C). These modeling findings are consistent with classic isotope-tracer studies in metazoans, which used radiolabeled or stable-isotope glucose to estimate PPP contribution to glucose metabolism and revealed tissue- and context-dependent differences in glucose utilization^7–12^. Our findings extend these isotope-tracer studies by moving from measurements in selected tissues to organism-wide metabolic-network modeling that predicts tissue-specific differences in preferred routes of glucose utilization.

A key finding from the whole-animal PPP model was the prediction, subsequently validated experimentally, that glucose-6-phosphate isomerase (GPI-1) can operate in the reverse direction *in vivo*^20,40^. This result is notable because GPI-1 sits at a metabolic junction between glycolysis and the PPP, reversibly catalyzing the interconversion of G6P and F6P. Given that G6P and F6P feed into the committed steps of the PPP and glycolysis, respectively, the directionality of GPI-1 flux could bias carbon flow between these pathways. To better understand how GPI-1 might shape differential glucose utilization across tissues, we next examined tissue-specific predictions of GPI-1 reaction directionality. For each cell type, we used the iMAT++ algorithm38 to infer the flux state of reactions (presence/absence and direction) by identifying an optimized network-wide flux distribution. These predictions indicated that GPI-1 exhibits relatively high reverse flux in reproductive tissues, including the germline and gonadal sheath cells (Fig. 1D)43. In contrast, tissues such as muscle and neurons displayed a different predicted GPI-1 flux profile, with less directional bias between PPP and glycolytic carbon flow (Fig 1D). Together, these modeling results predict tissue-specific differences in the capacity for PPP and glycolysis. Our modeling results also support a reverse GPI-1 flux preferentially favored in specific tissues, such as the germline. Importantly, these findings suggest that GPI-1 reaction directionality may help shape tissue-specific patterns of glucose utilization in metazoans.

### GPI-1 exhibits divergent, tissue-specific functions *in vivo*

To test whether the GPI-1–dependent metabolic programs predicted by our modeling contribute to *in vivo* physiology, we sought to genetically perturb GPI-1 function. To this end, we generated *C. elegans* strains carrying CRISPR–Cas9 knockouts of glucose-6-phosphate isomerase (*gpi-1*), phosphofructokinase (*pfk-1.1*) and glucose-6-phosphate dehydrogenase (*gspd-1*). While GPI-1 operates in both pathways via reversible isomerization of G6P to F6P, PFK-1.1 and GSPD-1 serve as rate-limiting enzymes that commit glucose flux toward glycolysis or the PPP, respectively (Fig. 2A). We reasoned that by benchmarking GPI-1 loss-of-function phenotypes against those of PFK-1.1 and GSPD-1, we could disentangle how GPI-1 contributes to anabolic versus catabolic metabolism *in vivo*. These genes in *C. elegans* share a high degree of conservation across evolution, including in mammalian species (Fig S1A - Fig S1D). Single-gene deletions were generated via removal of the whole coding sequences for *gpi-1, pfk-1.1* and *gspd-1* (Fig S1E) to provide a clean genetic deletion to examine the organismal consequences of disrupting key branch-point enzymes in glucose metabolism. Each of these enzymes is encoded by a single gene in *C. elegans,* save for a second gene encoding phosphofructokinase *(pfk-1.2)* that is expressed primarily in sperm (Fig S1F)^44^.

**Figure 2:**
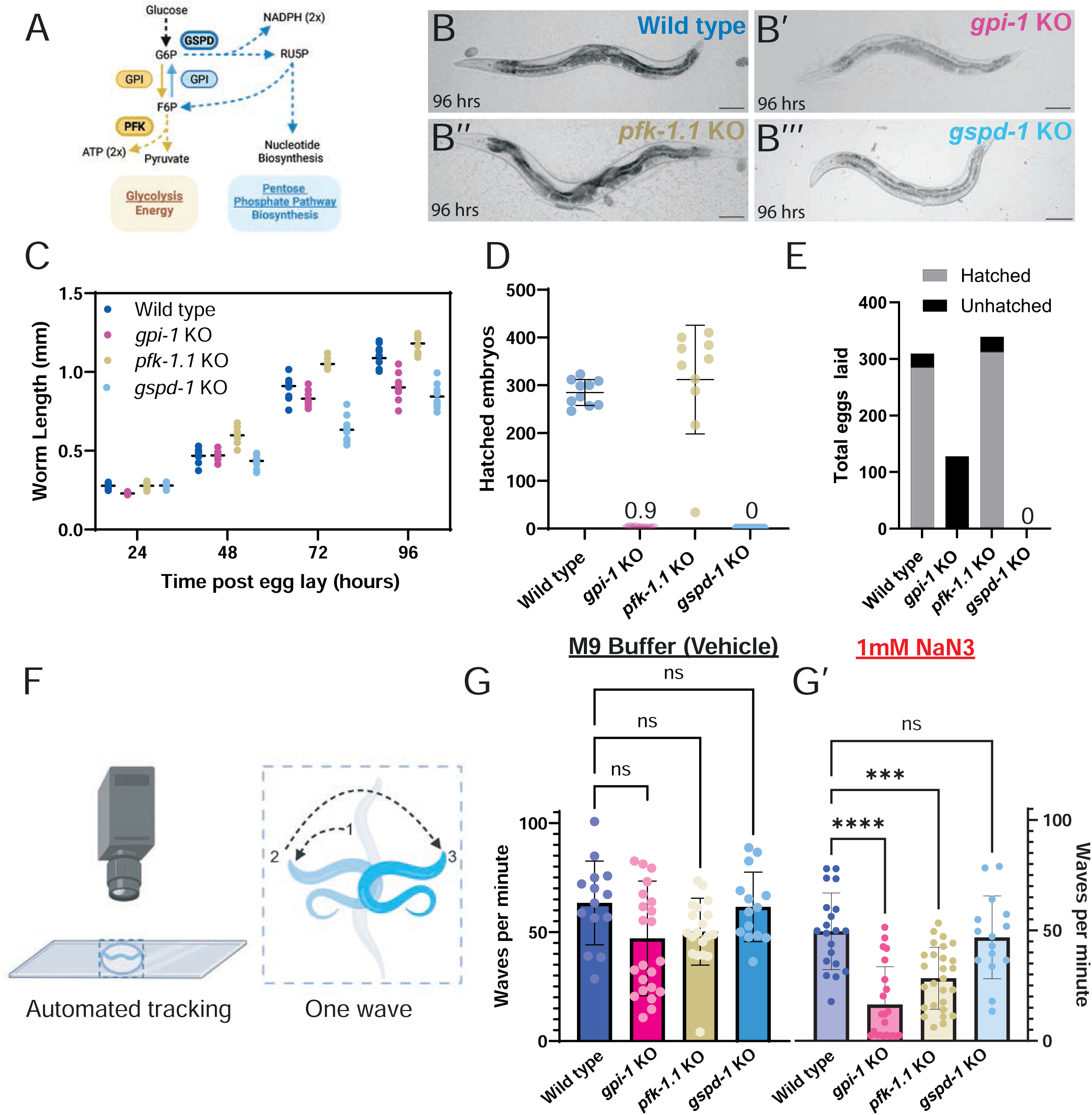
Glucose 6-phosphate isomerase (GPI-1) knockout results in impaired glycolysis and pentose phosphate pathway phenotypes. **(A)** Schematic of glycolysis and the pentose phosphate pathway enzymes PFK, GSPD and GPI and support PPP and glycolysis pathways. **(B)** Representative images of animals from wild type (B), *gpi-1* KO (B’) *pfk-1.1* KO (B’’) and *gspd-1* KO (B’’’) genotypes demonstrating different sizes at 96 hours post hatch. Scale Bar - 100 microns **(C)** Quantification of length for developmentally synchronized animals from wild type, *gpi-1* KO and *pfk-1.1* KO genotypes. **(D)** Quantification of hatched embryos from wild type, *gpi-1* KO, *pfk-1.1* KO and *gspd-1* KO animals displaying different reproductive fitness defects. **(E)** Total brood size quantification including hatched (gray) and unhatched (black) embryos across genotypes. **(F)** Diagram of swimming behavior analysis and quantification of thrashing in waves per minute **(G)** Swimming behavior quantification across genotypes in M9 vehicle buffer (G) and 1mM mitochondrial inhibitor sodium azide NaN_3_ (G’).

We then assessed several parameters of whole-animal physiology and metabolism that are underpinned by PPP and glycolytic metabolism. To assess physiological PPP function, we assayed larval development and reproductive fitness of adult animals. To assess glycolysis function, we assayed swimming behavior and quantitative readouts of metabolites via whole animal metabolomic changes and *in vivo* biosensors. These measurements reflect physiologically relevant biosynthetic demands as well as catabolic energy utilization and allowed us to systematically compare the physiological roles of glycolysis and the PPP within a single, tractable organism.

We started by quantifying body size during development and brood size across our knockout strains. *gpi-1* KO animals exhibited slow growth and reached the L4 stage later than wild type animals (Fig. 2B,C). We also observed that *gpi-1 (ola516)* mutants (hereafter referred to as *gpi-1* KO animals) exhibited severe embryonic lethality, producing fewer hatched and total embryos than wild type animals (Fig. 2D,E). In contrast, *pfk-1.1 (ola458)* mutants (hereafter referred to as *pfk-1.1* KO animals) displayed normal brood sizes (Fig. 2D,E) and, under our growth conditions (see Methods), grew at a similar rate to wild-type animals (Fig. 2B,C). *gspd-1 (ola594)* mutants (herein referred to as *gspd-1* KO animals) also exhibited severe sterility phenotypes and slow growth (Fig. 2B-2E), consistent with previous gene knockdown results45–47. These findings suggest that *gpi-1* and *gspd-1* share overlapping roles in reproductive fitness, consistent with our modeling predictions which indicated that GPI-1-dependent PPP flux is enriched in the germline.

Our modeling results indicated that GPI-1 and the reactions of glycolysis are enriched in muscles and neurons relative to other tissues (Fig 1C). To test whether GPI-1 functions in a canonical glycolysis role *in vivo,* we assayed animal swimming behavior. Swimming has been used as a paradigm for exercise^48,49^, as its sustained movement is predicted to require continuous ATP generation through glycolysis and oxidative phosphorylation. Therefore, we first used swimming behavior as a sensitive readout of coordinated somatic energy metabolism between tissues such as neurons and muscles. We quantified the swimming behavior^50,51^ (Fig. 2F) of *gpi-1, pfk-1.1* and *gspd-1* knockout (KO) animals in either M9 buffer (control) or in the presence of sodium azide (NaN₃), which inhibits mitochondrial respiration (Fig. 2G). We previously established that a loss-of-function allele for *pfk-1.1 (pfk-1.1(ola72)* only showed differences in swimming behavior compared to wild type while in the presence of NaN₃, suggesting that PFK-1.1 and glycolysis compensates during mitochondrial impairment^52^. *pfk-1.1* KO animals exhibited thrashing frequencies (waves per minute) comparable to wild type in M9 buffer (Fig. 2G), consistent with previous findings and the known compensatory roles of oxidative phosphorylation in *pfk-1.1* mutants^52^. However, under NaN₃ treatment, *pfk-1.1* KO animals showed significantly reduced thrashing, indicating that PFK-1.1–mediated glycolysis is required to sustain locomotor activity when oxidative phosphorylation is compromised (Fig. 2G′). In contrast, *gspd-1* KO animals showed no significant difference in thrashing relative to wild type in either condition, suggesting that PPP activity is dispensable for somatic energy metabolism during swimming (Fig. 2G,G′). *gpi-1* KO animals phenocopied the *pfk-1.1* KO animals in swimming behavior, displaying impaired swimming behavior compared to wild type animals treated with NaN₃ (Fig 2G’). The similarity between *gpi-1* KO and *pfk-1.1* KO swimming phenotypes suggests that GPI contributes to glycolytic energy production required for sustained locomotion and is consistent with a canonical function in glycolysis.

While it is currently not methodologically possible to perform tissue-specific metabolomics in *C. elegans* to confirm this dual, tissue-dependent role of GPI-1, we hypothesized that the observed metabolomic analysis of the whole animal would demonstrate changes in both glycolysis and PPP metabolites. To test this, we performed targeted metabolomics on *pfk-1.1* and *gpi-1* KO animals and compared glycolysis and PPP intermediates. Consistent with their roles in supporting somatic glycolysis, both mutants showed depletion of glycolytic metabolites relative to wild type animals, including fructose 1,6-bisphosphate (FBP) and lactate (Fig. S2A,B). Interestingly, *pfk-1.1* KO animals uniquely accumulated ribose 5-phosphate, a non-oxidative PPP intermediate (Fig. S2B). *gpi-1* KO animals showed depletion of reduced glutathione (GSH) relative to wild type animals (Fig. S2A), suggestive of a change in redox state. PPP is a major producer of NADPH which is necessary to convert oxidized glutathione (GSSG) to GSH for subsequent detoxification of reactive oxygen species. Direct comparison of the two mutants showed that, relative to *pfk-1.1* KO animals, *gpi-1* KO animals had significantly depleted PPP metabolites and GSH (Fig. 3A). Pathway analyses revealed similar depletion of glycolysis in both mutants, but opposite regulation of PPP metabolites (Fig. 3B). *gpi-1* and *pfk-1.1* knockout animals showed similar trends in differential PPP regulation when compared to wild type animals (Fig S2C,D). Together, these findings indicate that while *gpi-1* and *pfk-1.1* are both required to sustain glycolysis *in vivo*, GPI additionally supports PPP-dependent redox and biosynthetic demands, positioning GPI as a context-dependent branch-point controller of carbon allocation at the organismal scale.

**Figure 3:**
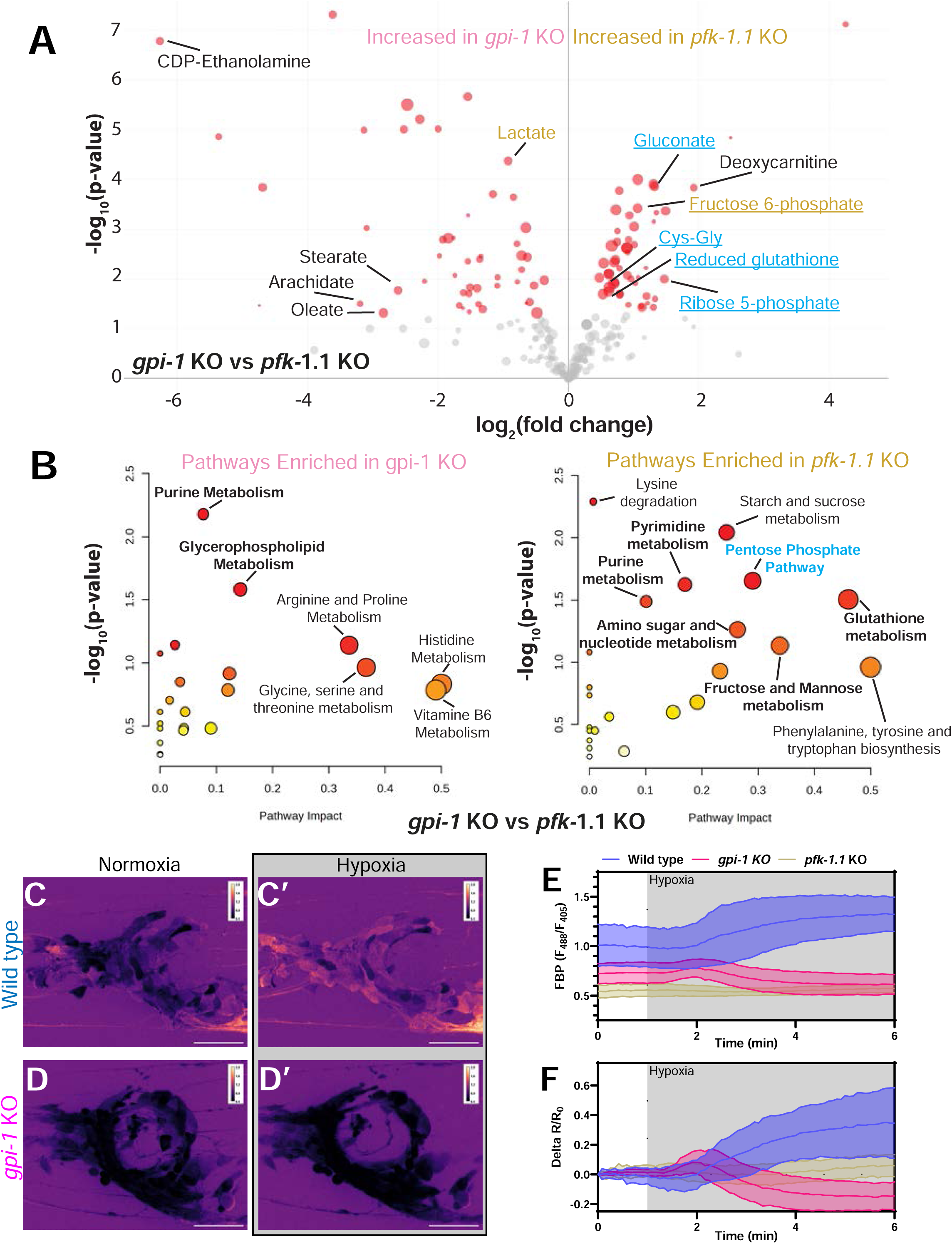
*gpi-1* KO animals exhibit depleted PPP and glycolytic metabolites. **(A)**Volcano plot for differential metabolites from targeted metabolomics comparing *gpi-1* KO animals and *pfk-1.1* KO animals. Red circles correspond to significantly differential metabolites, the size of the circle corresponds to abundance. Blue lettering indicates metabolites represented in either pentose phosphate pathway or redox metabolism. Gold lettering indicates glycolytic metabolite. Underlined metabolites are those that are differentially regulated in *pfk-1.1* or *gpi-1* KO animals relative to wild type controls. **(B)** Pathway analysis of significantly differential metabolites between *gpi-1* and *pfk-1.1* KO animals demonstrates affected pathways enriched in either mutant. Size of circle corresponds to pathway impact. Color corresponds to degree of significance (red - more significant, yellow - less significant). **(C-D)** Pan-neuronal expression of HYlight in wild type (C) and *gpi-1* KO animals (D) and responses to acute hypoxic stress (C’) and (D’). **(E-F)** Quantification of pan-neuronal HYlight responses demonstrating ratio over time (E) and change in ratio relative to the starting ratio (F).

To directly visualize glycolytic activity of GPI-1 *in vivo*, we monitored neuronal glycolysis using HYlight, a genetically encoded fluorescent biosensor that reports levels of the key glycolytic intermediate fructose-1,6-bisphosphate (FBP)^53^. HYlight enables real-time, cell-specific measurement of glycolytic flux in living animals, providing a quantitative readout of PFK-driven metabolism^15,41^. By coupling HYlight imaging with custom-built microfluidic devices, we can precisely control oxygen availability and mitochondrial function, allowing dynamic perturbation of cellular metabolism while continuously monitoring glycolytic responses^53,54^. We previously demonstrated that neuronal HYlight signals depend on *pfk-1.1* and increase in response to transient inhibition of mitochondrial function during hypoxic exposure, consistent with a role for glycolysis in meeting acute energy demands of neurons^15,41^.

Using this sensor, we examined how *gpi-1* contributes to both basal and stress-induced glycolysis. When HYlight was expressed under a pan-neuronal promoter, we observed that *gpi-1* KO animals were unable to sustain increases in glycolysis after hypoxia in all neurons, similar to *pfk-1.1* KO animals (Fig. 3C-F). This defect extended to the expression of HYlight under a single-cell promoter in the AIY interneuron—previously shown to rely on glycolysis to sustain synaptic function during hypoxic stress^52^—*gpi-1* knockout animals displayed normal baseline FBP levels but failed to elevate FBP following acute hypoxia (Fig. S2E-G). Together, these results demonstrate that *gpi-1* is required not only to maintain basal glycolytic activity in neurons, but also to dynamically upregulate glycolysis in response to mitochondrial inhibition. Genetically encoded sensors analogous to HYlight are not currently available to measure PPP activity *in vivo*, preventing us from directly testing the converse prediction in the reproductive tissues. Nevertheless, together with our genetic, physiological, modeling and metabolomic analyses, our data support a role for GPI-1 in somatic catabolism that appears distinct from its predicted anabolic functions in reproductive tissues.

### GPI-1 isoforms are differentially expressed in specific tissues *in vivo*

To determine the requirement of GPI-1 in specific tissues, we examined its endogenous expression in the intact animal. We generated a GPI-1::GFP knock-in at the *gpi-1* locus to visualize when and where endogenous GPI-1 is expressed *in vivo* (Fig. 4A). We observed broad somatic expression from embryos through adulthood (Fig. 4A-C, Fig. S3A - S3C). Beginning at larva stage 4 (L4), when animals reach reproductive maturity, GPI-1:GFP became prominent in the gonads (Fig 4D). Subcellular localization differed by tissue: GPI-1 was largely cytosolic in somatic tissues (Fig. 4C,4D’’) but showed pronounced subcellular enrichment in the developing germline and oocytes (Fig. 4D,4D’’’). Specifically, germline-restricted GPI-1 localized to membranous and vesicular structures within the rachis (Fig. 4D’), a shared cytoplasmic core of the syncytial germline that connects developing oocytes^56^. Subcellular enrichment to membranous structures was also evident in some embryos (Fig. S3D - S3E). Together, our findings indicate that GPI-1 is broadly expressed across tissues in the living animal, consistent with its broad roles in glucose metabolism. Our findings also reveal that GPI subcellular localization varies across tissues, with cytosolic distribution in somatic tissues, and germline-specific endomembrane compartmentalization. Interestingly, these differences in localization mirrored the organismal phenotypes we had observed for soma versus gonads in the *gpi-1-*associated phenotypes in catabolism versus anabolism.

**Figure 4:**
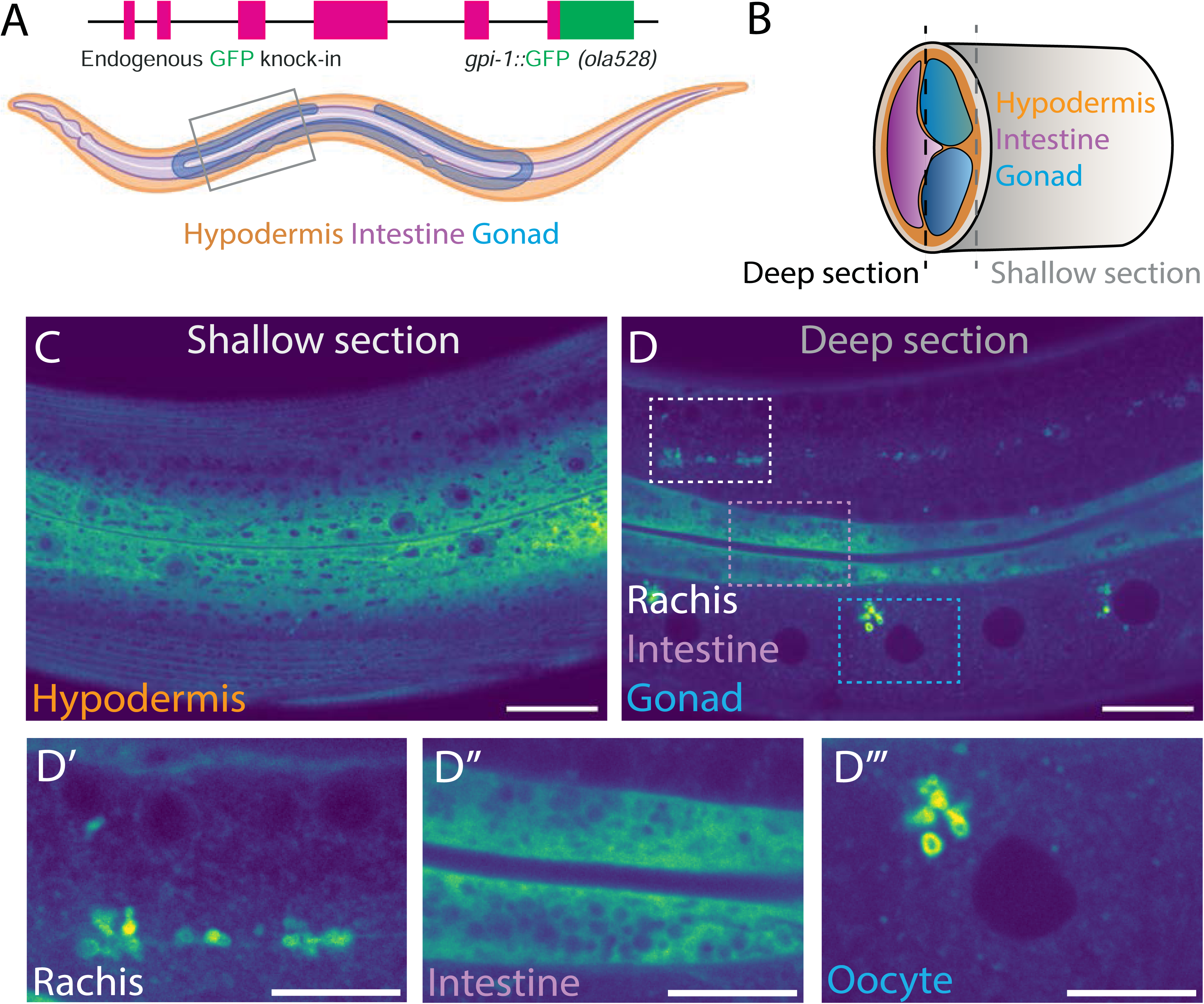
Endogenous labelling of *gpi-1* gene locus reveals differential localization patterns across tissues. **(A)** Schematic of GFP knock-in strategy at the genomic locus of *gpi-1* and *C. elegans* schematic with hypodermis, intestine and gonad of *C. elegans* hermaphrodites. **(B)** Schematic of transverse slice of the animal illustrating different sections examined to determine endogenous localization and expression pattern of *gpi-1 gene.* **(C)** Image from the **“**shallow section” of the animal demonstrating GPI-1::GFP expression in the hypodermis. Scale bar - 20 microns. **(D)** Image from the **“**deep section” of the animals demonstrating GPI-1B::GFP expression in the intestine and gonad. (D’ - D’’’) demonstrates the localization pattern of GPI-1::GFP in the gonadal rachis, intestine and oocytes respectively. (D) Scale bar - 20 micros. (D’ - D’’’) Scale bar - 10 microns.

To understand the molecular basis of GPI’s distinct localization patterns across tissues, we examined the *gpi-1* locus. The locus encodes two nearly identical protein-coding isoforms, GPI-1A and GPI-1B, which differ only by a 35 amino acid N-terminal domain present in GPI-1B (Fig. 5A,B). Therefore, we engineered GFP knock-in alleles that selectively label each isoform in its native genomic context to investigate whether these two isoforms localize differently. This was achieved by disrupting the alternative start site of one isoform while preserving expression of the other, allowing us to visualize each protein independently under endogenous regulatory control (Fig, 5A,B, lower right).

**Figure 5:**
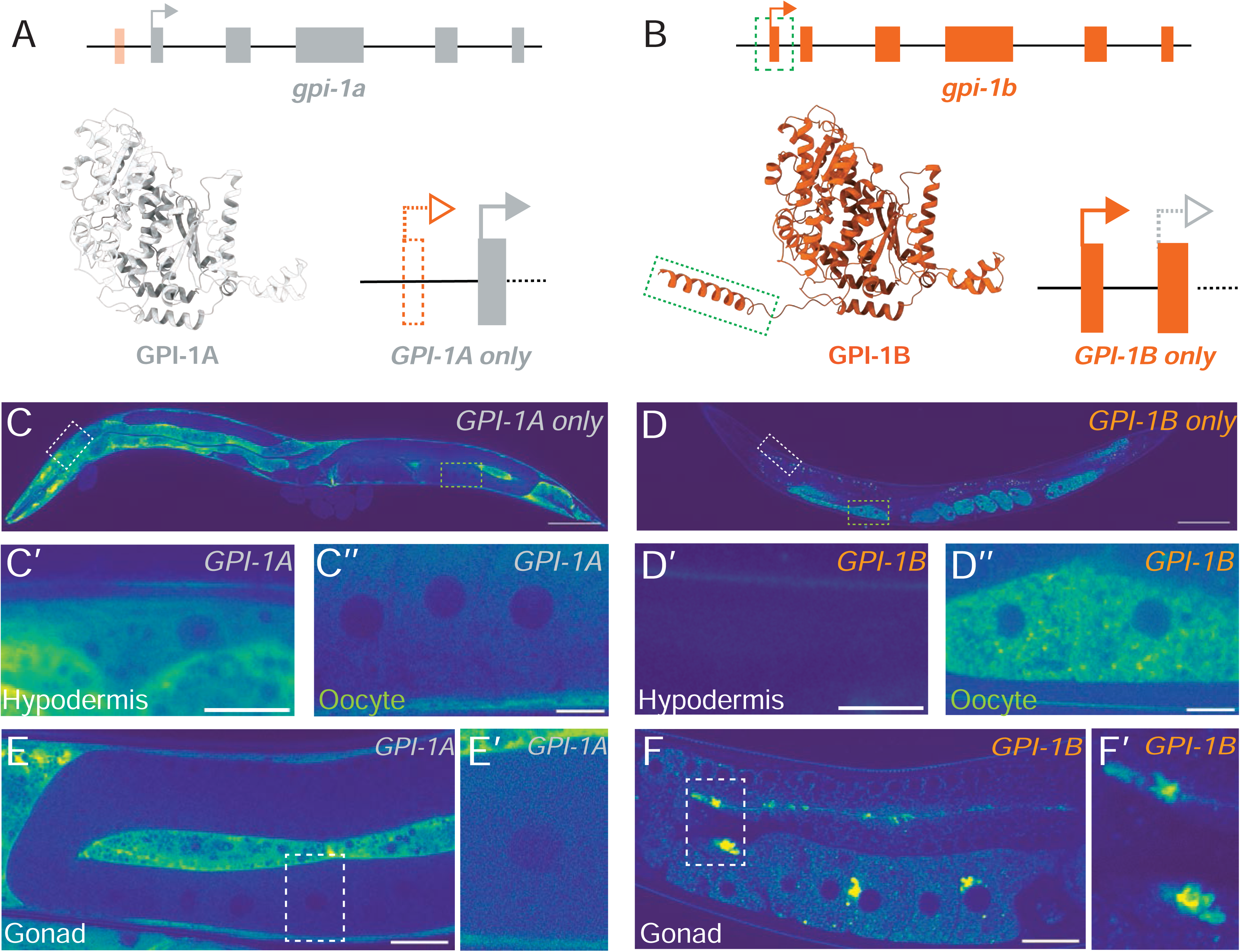
GPI-1 isoforms exhibit distinct patterns of tissue expression and subcellular localization. **(A)** (Top) Schematic of *gpi-1* gene encoding for GPI-1A isoform in gray. (Lower left) predicted AlphaFold structure of GPI-1A protein. (Lower right) design of *GPI-1A only* mutant isolating GPI-1A expression by removing upstream exon encoding the GPI-1B isoform (dashed orange box, also orange box in schematic of gene on top). **(B)** (Top) Schematic of *gpi-1* gene encoding for GPI-1B isoform in orange. (Lower left) predicted AlphaFold structure of GPI-1B protein. (Lower right) design of *GPI-1B only* mutant isolating GPI-1B expression by mutagenizing GPI-1A start codon. Green box indicates exon specific to GPI-1B which encodes GPI-1B N-terminal protein domain also marked in green. The dashed start site on the second exon represents the mutated start site for GPI-1A in this strain (see Methods) **(C)** *GPI-1A only* animals demonstrate broad somatic tissue expression and cytosolic localization in the hypodermis (C’) and in oocytes (C’’) **(D)** *GPI-1B only* animals demonstrating sparse somatic tissue expression and subcellular localization in the hypodermis (D’) and in oocytes (D’’). (C and D) Scale bar - 100 microns (C’, C’’, D’ and D’’) Scale bar - 10 microns. **(E)** GPI-1A expression in the adult gonad with zoom-in image of subcellular GPI-1 localization in an oocyte (E’). Scale Bar - 20 microns **(F)** GPI-1B expression in the adult germline with zoom-in image of subcellular GPI-1 localization in an oocyte and in the ovary (F’). Scale Bar - 20 microns.

In the GPI-1A–expressing strain (*GPI-1A only* in figures), we observed broad tissue expression of GPI-1A::GFP from embryos through adulthood (Fig. S3F, Fig. 5C). The localization of GPI-1A::GFP was largely cytosolic in somatic tissues (Fig. 5C’) and the subcellular germline-specific foci characteristic of the wild-type *gpi-1::GFP* background were absent (Fig. 5C’’ and Fig. 5E). In contrast, the GPI-1B–expressing strain (*GPI-1B only* in figures) displayed low levels of somatic expression starting from the L1 larval stage through adulthood (Fig. S3G, Fig. 5D’) and pronounced germline-localized endomembrane localization starting at the L4 and young adult stage (Fig. 5D’’ and Fig. 5F), consistent with the germline-specific structures observed in the wild-type *gpi-1::GFP* strain (Fig. 4D). These localization patterns were also observed in laid embryos (Fig. S3H-J).

Together, these findings suggest that the distinct localization patterns observed in the endogenous *gpi-1::gfp* strain arise from the differential expression and localization of the GPI-1A and GPI-1B isoforms across somatic and germline tissues.

### GPI-1B localizes to the endoplasmic reticulum via its N-terminal helical domain

Our observations suggest that GPI-1 function is spatially partitioned, and that different isoforms are differentially expressed in those tissues. We thus set out to elucidate the site of GPI-1 localization. We started by overexpressing each isoform’s cDNA translationally fused to GFP in the same defined neurons—the AIY interneuron (using the cell-specific *pttx-3* promoter) and the GABAergic VD/DD neurons (using the cell-specific *punc-47* promoter) in wild type animals (Fig 6A). We observed that GPI-1A::GFP was diffusely cytosolic throughout the soma and neurites in both neuronal types (Fig. 6B; Fig. S4A), consistent with the observed cytosolic localization when it was expressed from the endogenous locus. In contrast, GPI-1B::GFP was enriched at subcellular compartments (Fig. 6B’; Fig. S4A’), mirroring the punctate, endomembrane localization observed in the germline. Together, these results indicate that the two isoforms have intrinsically distinct subcellular localization patterns, independent of the cell type in which they are expressed.

**Figure 6:**
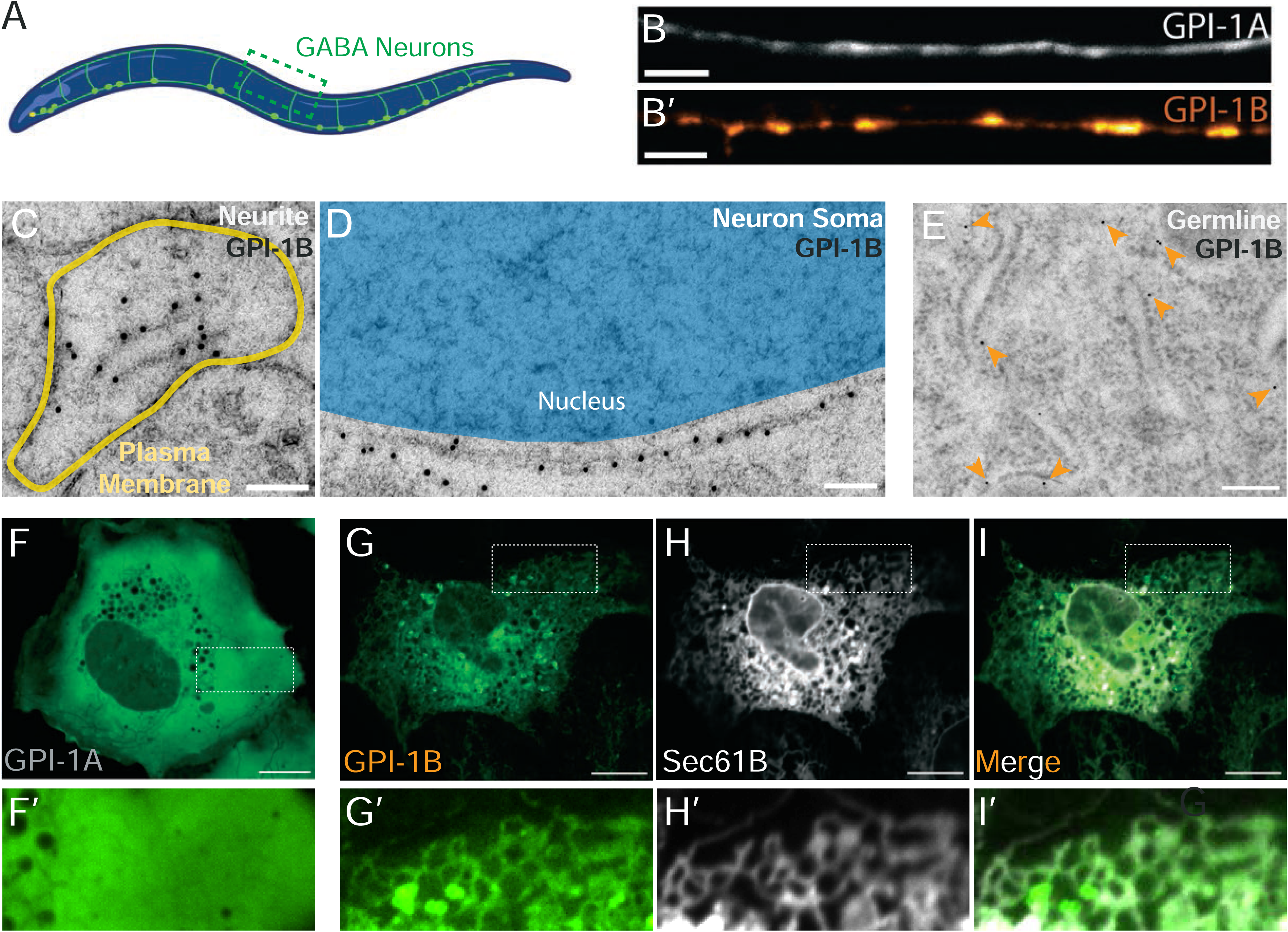
GPI-1B localizes to the endoplasmic reticulum. **(A)** Schematic of *C. elegans* GABAergic neurons. **(B)** Exogenous cDNA expression of GPI-1A (B) and GPI-1B (B’) in GABAergic neurons demonstrating localization pattern differences between GPI-1 isoforms. Scale bar - 5 microns. **(C)** Immuno-EM of *C. elegans* wild type with pan-neuronal GPI-1B::GFP expression showing GPI-1B signal in neurites near ER membrane. Yellow border denotes neurite plasma membrane, Scale bar - 100nm. **(D)** Immuno-EM of *C. elegans* wild type with pan-neuronal GPI-1B::GFP expression showing GPI-1B signal in neuron soma near ER membrane. Blue region denotes nucleus, Scale bar - 100nm. **(E)** Immuno-EM of *C. elegans GPI-1B only* strain showing GPI-1B signal (orange arrows) in the germline near ER membranes. Scale bar 100nm. **(F)** Expression of GPI-1A in RPE-1 cells with inset image (F’) showing cytosolic localization pattern. **(G)** Expression of GPI-1B in RPE-1 cells with inset image (G’) showing subcellular, membranous localization pattern. **(H)** Co-expression of general ER marker Sec61B in RPE-1 cells with inset image (H’) **(I)** Merge image showing colocalization of GPI-1B and general ER marker Sec61B in RPE-1 cells with inset image (I’) (F-I) Scale bar - 10 microns.

To determine the precise subcellular localization of GPI-1B in *C. elegans*, we performed immuno–electron microscopy (EM) on animals pan-neuronally overexpressing GPI-1B (using the pan-neuronal *prab-3* promoter) and GPI-1B only animals. In the pan-neuronal strain overexpressing GPI-1B, signal was detected on membranes within neurites and near the nucleus within neuronal somas (Fig. 6C, 6D). To further examine subcellular localization *in vivo*, we generated a single-copy insertion strain expressing GPI-1B::GFP in the DA9 motor neuron (by using the cell-specific *pitr-1* promoter). GPI-1B localized to axons, dendrites, and the DA9 soma (Fig. S4B), where Airyscan super-resolution confirmed perinuclear and reticular structures characteristic of ER membranes (Fig. S4C). Colocalization analysis of DA9 cell somas also revealed GPI-1B localization which resembled that of the ER marker SP12 (Fig. S4D). In the rachis of *gpi-1b::gfp* animals, immuno-EM revealed GPI-1B signals adjacent to ribosome-rich regions, consistent with ER or ER-associated structures (Fig. 6E). Together, these findings suggest that GPI-1B localizes to the ER or ER-associated endomembrane structures in *C. elegans*.

We sought to further support our observations of GPI-1B ER localization by examining isoform localization in a heterologous single-cell system. Retinal pigment epithelium (RPE-1) cells provide a simplified cellular context relative to the intact worm and contain well-defined organelles that allow unambiguous identification of the subcellular localization of the GPI-1 isoforms. We therefore expressed the *C. elegans* GPI-1 isoforms in RPE-1 cells and noted their subcellular localization. Consistent with our observations in *C. elegans* neurons, we found that GPI-1A displayed diffuse cytosolic localization in RPE-1 cells, whereas GPI-1B accumulated in endomembranous structures throughout the cytoplasm (Fig. 6F,G). Co-labeling of GPI-1B and the ER marker Sec61β revealed strong colocalization (Fig. 6G-6I). In some cells we also observed GPI-1B enrichment on ER-derived lipid droplets (Fig. S4E–G). While we did not test vertebrate GPI isoforms, the endomembrane localization of GPI-1B in cells of different animals, and the absence of such targeting for GPI-1A, suggests that the two isoforms carry distinct intrinsic signals that may underlie divergent but conserved targeting mechanisms. Importantly, our findings suggest distinct localization of these isoforms due to protein-specific properties.

We next asked which structural features of GPI-1B enable its subcellular localization. GPI-1A and GPI-1B are identical except for a 35–amino acid N-terminal domain (NTD) encoded by the GPI-1B–specific 5′ exon (Fig. 5 A,B). We therefore hypothesized that this NTD functions as a localization sequence. AlphaFold structure predictions (Fig. 7A) suggest that the NTD forms a helical element containing residues consistent with amphipathic or coiled-coil properties. To test whether this region is required for localization, we introduced point mutations into the NTD predicted to disrupt these motifs (Fig. 7B). The resulting coiled-coil mutant (CCM) GPI-1B::GFP no longer formed punctate structures and instead localized diffusely in the DA9 motor neuron, similar to GPI-1A (Fig. 7C,D′). To test this requirement *in vivo* at endogenous expression levels, we used CRISPR to introduce the CCM allele into the *gpi-1::gfp* background. In contrast to wild-type GPI-1B, which forms discrete germline foci, GPI-1(CCM)::GFP was diffusely cytosolic (Fig. 7D,E). Expression of the CCM variant in RPE-1 cells yielded the same result: the mutant protein was cytosolic and failed to localize to ER-associated lipid droplets (Fig. 7F,F’; S4H–J). Together, these data demonstrate that the GPI-1B N-terminal helical domain is necessary for subcellular localization to ER membranes, and ER-associated endomembrane structures, in tissue culture cells and *in vivo*. These findings are consistent with other observations of small sequence variations in metabolic enzymes regulating subcellular targeting, thereby influencing metabolic output through spatial organization of enzymes^21,57–59^.

**Figure 7:**
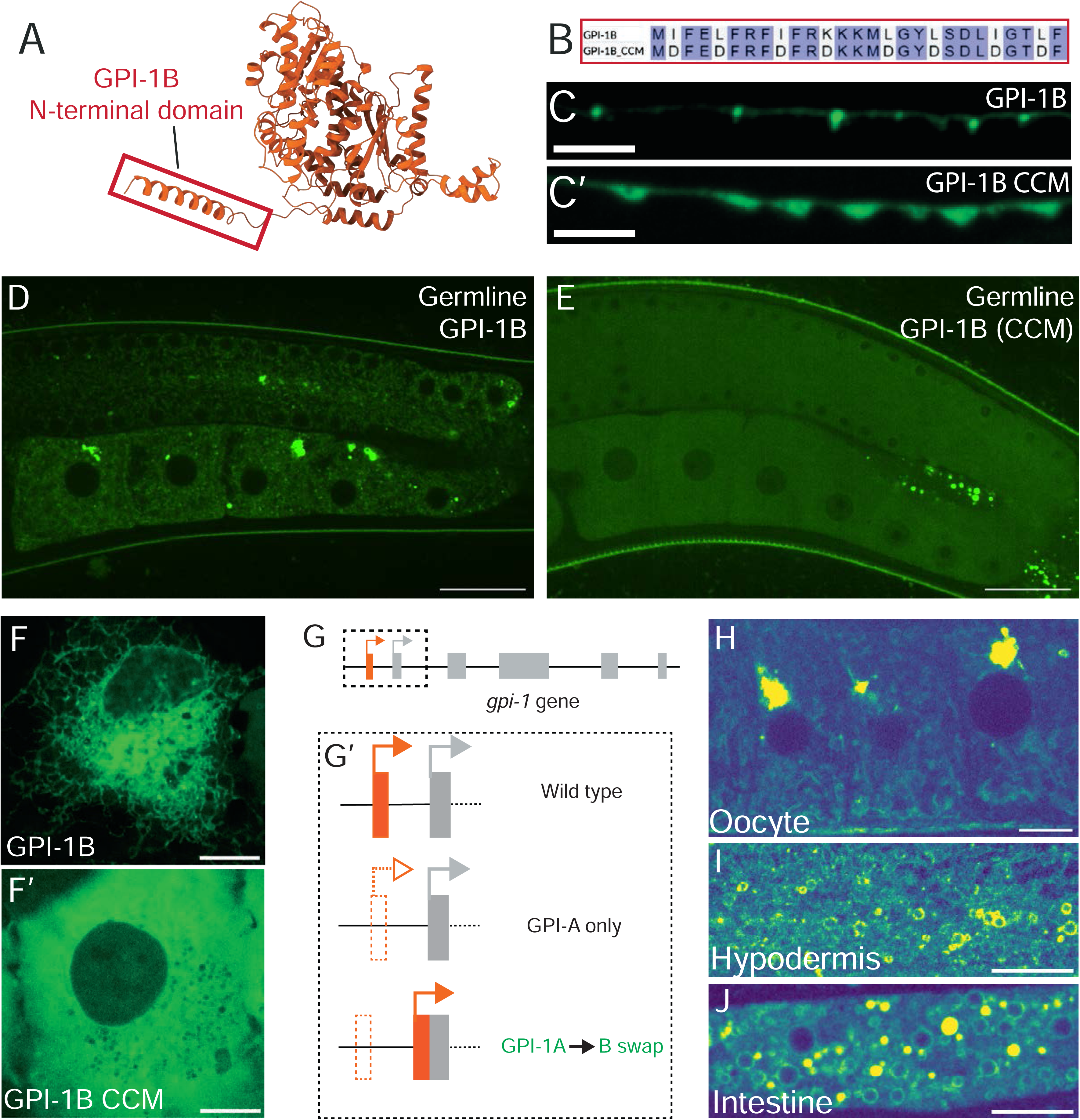
GPI-1B localizes via its N-terminal helical domain. **(A)** Predicted AlphaFold structure for *C. elegans* GPI-1B, with GPI-1B-specific N-terminal domain (NTD) outlined in red. **(B)** Alignment of wild type GPI-1B NTD in red and recombinant coiled-coil mutant (CCM) NTD below illustrating NTD mutagenesis **(C)** Subcellular localization pattern of wild type GPI-1B (C) and CCM GPI-1B (C’) tagged with GFP in DA9 neuron axons showing difference in subcellular localization. Scale bar - 5 microns **(D)** *GPI-1B only* animals showing subcellular localization of GPI-1B in the germline **(E)** *GPI-1B (CCM) only* animals showing cytosolic localization of GPI-1B in the germline. Scale bar - 20 microns. **(F)** GPI-1B expression in RPE-1 cells showing subcellular localization (F’) GPI-1B CCM expression demonstrating cytosolic localization. Scale bar - 10 microns. **(G)** Schematic of *gpi-1* gene locus with GPI-1A (gray) and GPI-1B (orange) coding isoforms. **(G’)** Schematic of putative GPI-1B swap strain constructed by first removing the GPI-1B-specific exon followed by reinsertion of the GPI-1B exon upstream of GPI-1A-specific translation start site. **(H)** Oocyte in GPI-1B swap animal showing subcellular GPI-1 localization. **(I)** Hypodermis in GPI-1B swap animal displaying subcellular GPI-1 localization **(J)** Intestine in GPI-1B swap animal displaying subcellular GPI-1 localization

Given that the major differences between GPI-1A and GPI-1B appear to be their tissue expression and subcellular localization (Fig. 6,7), we next asked whether the GPI-1B-specific domain is sufficient for subcellular localization *in vivo*. We thus engineered a strain that expresses GPI-1B in place of GPI-1A throughout the organism. Using CRISPR, we fused the GPI-1B-specific 5’ exon into the GPI-1A–only strain, thereby converting all endogenous GPI-1A expression into GPI-1B expression (hereafter referred to as the “GPI-1B swap” strain; Fig. 7G). We observed that in the GPI-1B swap strain, GPI-1 no longer displayed diffuse cytosolic localization in somatic tissues. Instead, it formed prominent membranous and vesicular foci in somatic tissues such as the hypodermis and intestine (Fig. 7I,J). This strain also restored the germline localization that was absent in the GPI-1A–expressing strain (Fig. 7H). These findings confirm that subcellular compartmentalization of GPI-1B is an intrinsic property of the protein and does not depend on the tissue in which it is expressed. Given our observations regarding GPI’s distinct physiological roles in these tissues, our findings also suggest that these isoform-specific differences may reflect, or even contribute to, context-dependent regulation of catabolic or anabolic metabolism *in vivo*.

### GPI-1 isoforms differentially contribute to tissue-specific functions

We next sought to understand whether and how isoform expression and localization contribute to GPI-1 function. We examined this by selectively deleting each *gpi-1* isoform via the engineering of strains that express either GPI-1A or GPI-1B from the endogenous locus. To then determine whether the isoforms also differentially support somatic glycolysis, we analyzed swimming behavior in the single-isoform mutants, a phenotype impaired in the full *gpi-1* knockout. GPI-1A–expressing animals (lacking GPI-1B) thrash at levels comparable to wild type in both M9 buffer and NaN₃ (Fig. 8A,8A′). In contrast, GPI-1B–expressing animals (lacking GPI-1A) displayed significantly reduced thrashing even in M9, with further impairment in NaN₃ (Fig. 8A and 8A′). These results show that GPI-1A is necessary and sufficient to support swimming behavior, and suggest a role for the GPI-1A isoform in somatic glycolytic activity required for sustained movement.

**Figure 8:**
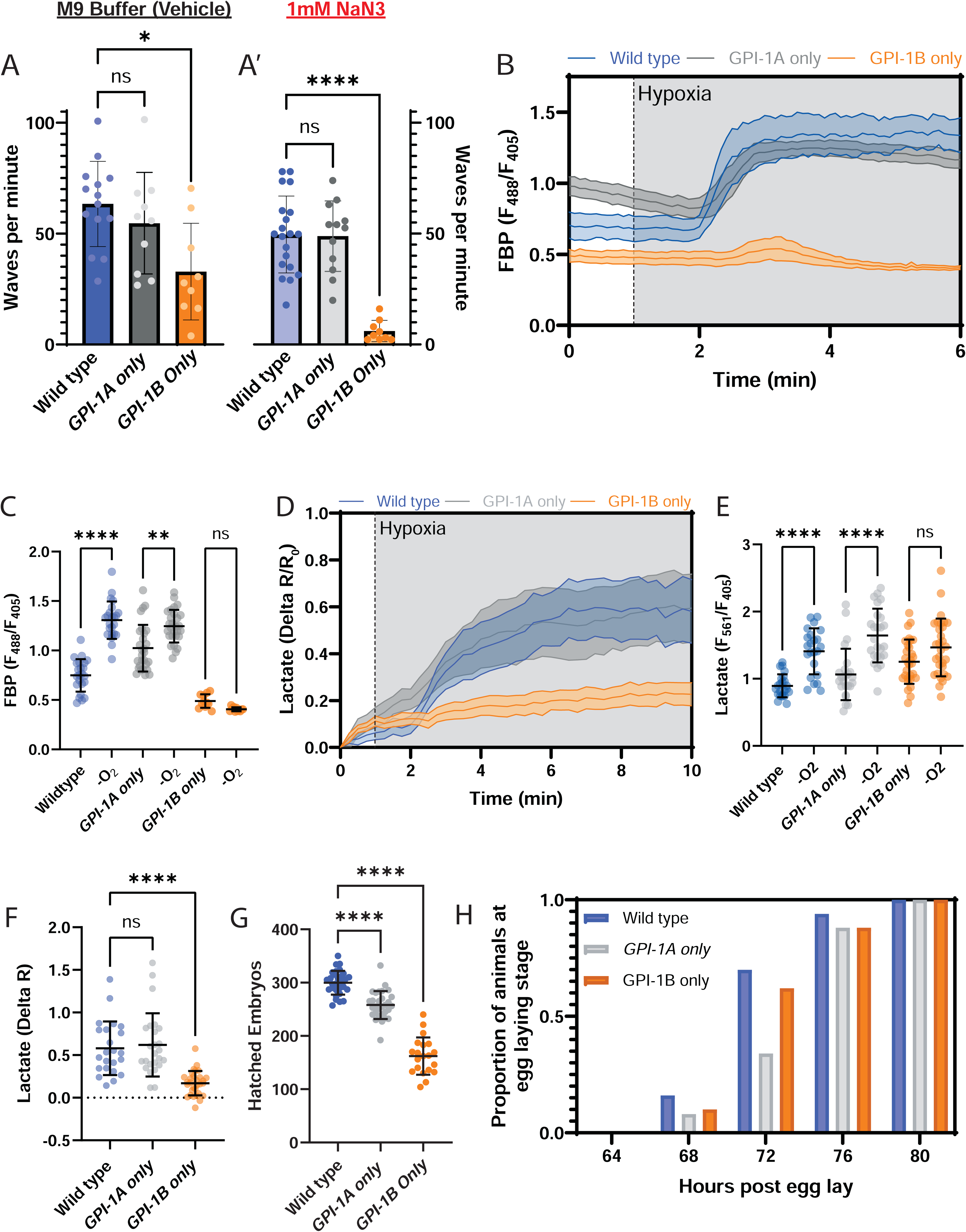
GPI-1 isoforms differentially regulate reproductive fitness and somatic glycolysis across tissues. **(A)** Swimming assay quantification for single isoform mutants in the M9 vehicle buffer condition (A) and 1mM mitochondrial inhibitor NaN3 condition (A’). **(B)** AIY HYlight ratiometric measurements over time of single isoform mutants in normoxia (white) and hypoxia (gray). **(C)** Baseline and hypoxic (-O_2_) values for AIY HYlight in single isoform mutants. **(D)** Relative ratio change over time (Delta R/R_0_) of AIY FiLa lactate biosensor in response to acute hypoxia (gray) in single isoform mutants **(E)** Baseline and hypoxic (-O_2_) values for AIY FiLa in single isoform mutants. **(F)** Delta R value representing final relative change in FiLa Lactate ratio over time in response to acute hypoxia. **(G)** Quantification of hatched embryos over total reproductive lifespan in single isoform mutants compared to wild type animals. **(H)** Proportion of age-synchronized animals at egg laying stage at different time points post hatching in single isoform mutants.

To directly assess how each isoform contributes to somatic glycolysis, we then examined neuronal FBP dynamics using the HYlight biosensor. In the full *gpi-1* knockout, neuronal HYlight signals are reduced and fail to increase following mitochondrial inhibition upon transient hypoxia, indicating a defect in glycolytic regulation (Fig. 2A). GPI-1A–expressing animals with cell-specific expression of HYlight in the AIY interneuron demonstrated increases in FBP following acute hypoxia (Fig. 8B,8C). In contrast, HYlight in GPI-1B–expressing animals failed to upregulate glycolysis upon transient hypoxia (Fig. 8B,8C). These data suggest that GPI-1A, but not GPI-1B, supports glycolytic flux in somatic tissues like neurons.

To independently validate these findings, we imaged lactate levels using FiLa, a genetically encoded fluorescent biosensor that reports intracellular lactate production^60,61^. Lactate is a direct downstream product of glycolysis and measuring lactate provides a complementary metabolic readout to measuring FBP via HYlight, enabling confirmation of glycolytic flux in the examined genetic backgrounds. Consistent with HYlight measurements, wild type and GPI-1A–expressing animals increased lactate production during transient hypoxia, whereas GPI-1B–expressing animals did not (Fig. 8D-F). The concordance between two orthogonal metabolic reporters, FBP and lactate, imaged *in vivo* provides strong evidence that GPI-1A is necessary and sufficient to drive somatic glycolysis, while the presence of the GPI-1B gene locus alone is insufficient to support this function.

Unlike glycolysis, there are no *in vivo* sensors to directly examine PPP. We therefore assayed the effects of individual isoforms on reproductive fitness, motivated by our previous findings of germline PPP enrichment and dependence. Given the severe reproductive defects of the full *gpi-1* knockout, we also quantified brood sizes in the single-isoform mutants. Neither strain recapitulated the embryonic lethality and sterility of the complete knockout (Fig. 2D,E), indicating partial functional redundancy of the two isoforms. However, we observed that both GPI-1A–expressing animals and GPI-1B–expressing animals produced significantly fewer hatched embryos than wild type over the course of their reproductive lifespan (Fig. 8G), suggesting that both isoforms are necessary in the germline to recapitulate the wild type phenotype. Interestingly, GPI-1A–expressing animals (which lack GPI-1B) showed delayed egg production: at 72 hours post-hatching, a smaller fraction contained embryos compared to wild type and GPI-1B-expressing animals (Fig. 8H). Together, these results indicate that the two isoforms act in partially redundant but functionally specialized roles, with GPI-1B, the isoform enriched in the germline, being specifically required for normal brood size and reproductive timing in the germline.

### GPI-1B can metabolically function in somatic tissues

Next we wanted to examine how changing GPI-1 localization affects metabolic and physiologic function. To address this we assayed the GPI-1B swap strain that forces GPI-1B expression in somatic tissues for somatic and germline phenotypes. We first observed that the GPI-1B swap strain displayed normal swimming behavior in both M9 and NaN₃ conditions (Fig. 9A,A′), suggesting that somatic glycolytic activity required for sustained movement can be supported by forcing expression of GPI-1B. These results are consistent with the partial redundancy of both isoforms that we observed in the germline, and suggest that the impaired swimming phenotype of GPI-1B-expressing animals arises at least in part from the low somatic expression of this isoform. This interpretation is consistent with transcriptomic data showing that GPI-1A is the predominantly expressed isoform in most examined neurons^62^.

**Figure 9:**
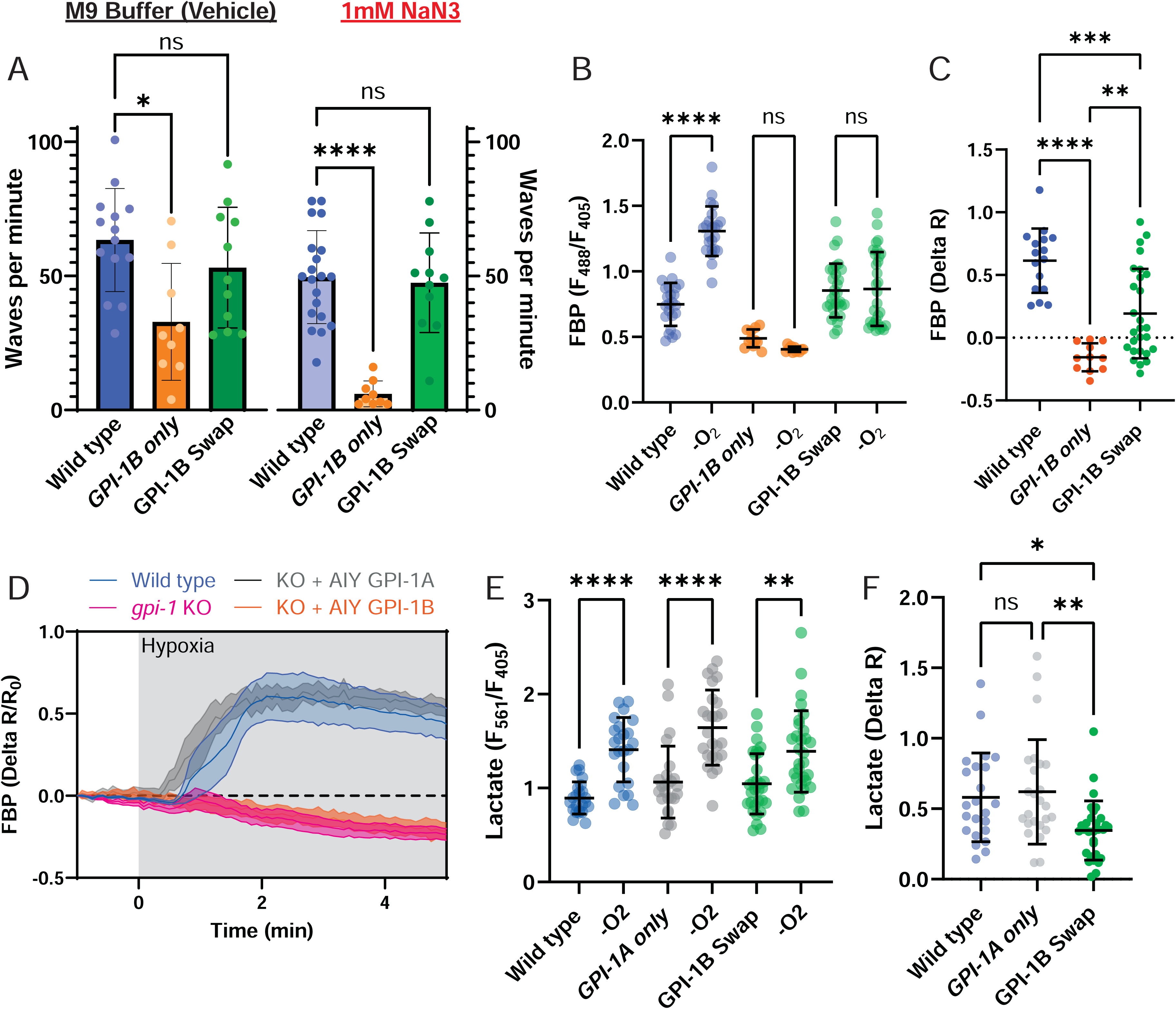
GPI-1B partially recapitulates functional glycolysis in somatic tissues. **(A)** Swimming assay quantification for *GPI-1B only animals* and GPI-1B swap animals compared to wild type animals in the M9 vehicle buffer condition and 1mM mitochondrial inhibitor NaN3 condition **(B)** Baseline and hypoxic (-O_2_) values for AIY HYlight in *GPI-1B only* and GPI-1B swap animals compared to wild type animals. **(C’)** Delta R value representing final relative change in HYlight FBP ratio over time in response to acute hypoxia. **(D)** Relative change in ratio over time (Delta R/R_0_) for wild type and *gpi-1* KO animals with AIY-specific rescue demonstrating differential rescue of HYlight responses to acute hypoxia. Wild type (blue) and *gpi-1* KO animals (pink). AIY-specific expression of GPI-1A (+AIY GPI-1A - gray) and GPI-1B (+AIY GPI-1B - orange) in the *gpi-1* KO background shows different patterns of rescue HYlight ratio. **(E)** Baseline and hypoxic (-O_2_) values for AIY Lactate in *GPI-1B only* and GPI-1B swap animals compared to wild type animals. **(F)** Change in FiLa ratio (Delta R) from normoxia to hypoxia for wild type, *GPI-1B only* animals, and GPI-1B swap animals.

To directly assess how GPI-1B supports glycolysis in somatic tissues, we monitored neuronal FBP and lactate dynamics in GPI-1B swap animals via the HYlight and FiLa biosensors, respectively. Interestingly, we observed that in the GPI-1B swap strain, FBP increases following transient hypoxia were significantly reduced, producing an intermediate phenotype between wild type and *GPI-1B only* animals (Fig. 9B,C) and failing to recapitulate the wild type-like phenotype observed for *GPI-1A only* animals (Fig 8B,C). We further supported these findings by measuring FBP responses to hypoxia in *gpi-1* KO animals with cell-specific expression of either GPI-1A or GPI-1B cDNA. Cell-specific expression of each isoform in AIY neurons yielded consistent results—GPI-1A expression restored hypoxia-induced increases in HYlight, while GPI-1B expression did not (Fig. 9D). Measurements of cellular lactate confirmed this deficit in glycolytic dynamics for *GPI-1B swap* animals: although lactate increased during hypoxia, the response amplitude of *GPI-1B swap* animals was significantly reduced relative to wild type or to *GPI-1A only* animals (Fig. 9E,F). These findings indicate that GPI-1B can partially compensate for GPI-1A to maintain basal glycolysis, but is insufficient to support the full dynamic regulation of glycolytic flux observed for wild type or *GPI-1A only* animals. These findings suggest that isoform function is partially redundant yet distinct, potentially reflecting regulation at the level of enzyme localization, consistent with differences in GPI-1 subcellular distribution.

Together, our data demonstrate that the two GPI-1 isoforms differ in their tissue expression, subcellular localization, and ability to support glucose metabolism *in vivo*. We observe that GPI-1A is broadly expressed in somatic tissues, localizes diffusely in the cytosol, and is sufficient to support glycolysis in somatic tissues, whereas GPI-1B is enriched in the germline, localizes to endomembrane compartments, and supports phenotypes associated with anabolism such as normal brood size and reproductive fitness, perhaps differentially supporting, via its localization, PPP reactions enriched within the germline of the animal.

## DISCUSSION

We show that GPI, a glycolytic and PPP enzyme that reversibly catalyzes the interconversion of G6P and F6P, supports distinct anabolic and catabolic functions *in vivo* via divergent, tissue-specific mechanisms of metabolic regulation. Metabolism is often conceptualized as a set of universally deployed biochemical pathways, yet tissues differentially use these conserved enzymes to fulfill distinct anabolic and catabolic demands. Using CRISPR knockouts, live metabolic imaging, and isoform-swap strains, we find that GPI-1A supports cytosolic glycolysis and rapid energetic responses in somatic tissues, whereas GPI-1B localizes to ER-associated membranes, supports germline metabolism, and requires a short N-terminal helical motif for targeting. These data suggest that metabolic control *in vivo* extends beyond enzyme abundance: differential distribution of metabolic isoforms across tissues may contribute to tissue-specific metabolic configurations, where cytosolic GPI-1A enables dynamic glycolytic flux in neurons and somatic cells, and both isoforms cooperate to support the biosynthetic and PPP flux of the germline. Together, this work provides a model for how a single enzyme can differentially support anabolic and catabolic programs across tissues, and establishes *C. elegans* as a tractable model to dissect compartmentalized metabolism *in vivo*.

Our findings reveal that GPI-1 is not functionally uniform *in vivo*, but instead performs distinct metabolic functions across tissues, with physiological consequences that only emerge at the level of the whole organism. Specifically, modeling of tissue-specific metabolism via analysis of single-cell RNA seq data revealed differential patterns of glucose metabolism, whereby GPI-1 demonstrated high flux across multiple tissues but for different pathways. While we focus on glycolysis and PPP to demonstrate the divergent usage of a particular branch point in glucose metabolism, this approach can be extrapolated to generate a comprehensive map of transcriptionally-favored metabolic reactions in the context of specific adult tissues. We complement these modeling approaches by then assessing tissue-specific physiology within whole gene knockouts. Although GPI-1 is broadly expressed, loss of *gpi-1* resulted in two separable phenotypes: defects in fertility and embryonic development, and failure of somatic tissues to sustain glycolysis under energetic stress. These outcomes cannot be inferred from single-cell or biochemical studies, which typically treat GPI as a generic glycolytic enzyme. Rather, in the intact animal, tissues differentially partition carbon flux, and the germline and soma experience fundamentally different metabolic pathway outcomes.

Disrupting GPI at the whole-organism level therefore uncovers tissue-specific susceptibilities underpinned by impaired anabolic investment in reproduction and catabolic energy production required for behavior and neuronal function. The fact that these phenotypes emerge only when metabolism is studied across interacting tissues underscores the importance of whole-animal approaches to understand how reversible metabolic nodes are regulated to support organismal fitness. Our findings are consistent with studies in mammals showing that in disease states in which glucose-6-phosphate dehydrogenase (G6PD) function is compromised, both humans and mice display growth defects, infertility, and sensitivity to oxidative stress^46,47,63^. We extend these findings by now demonstrating that knockouts of these enzymes in whole-animals result in different physiological phenotypes in specific tissues, extending our understanding of how anabolic and catabolic glucose utilization are differentially coupled to organismal fitness *in vivo*. Our study integrates current cutting-edge approaches to investigate tissue-specific metabolism, but it does not directly measure metabolite flow, particularly for PPP, across pathways and tissues. Our study integrates current cutting-edge approaches to dissect tissue-specific metabolism, yet comes short of direct measurement of metabolites across pathways and tissues. For example, genetically encoded biosensors that directly report PPP metabolites are not currently available; instead, the closest readouts of PPP activity come from metabolomic approaches, isotope-labeling experiments, or sensors that measure NADPH as a proxy for PPP activity. These limitations, together with our findings of tissue-specific roles for GPI-1, highlight the need to develop sensors and approaches that can directly probe specific metabolic pathways in defined tissues *in vivo*.

A longstanding assumption in metabolism is that glycolytic enzymes, being ancient, highly conserved, and biochemically similar across tissues, function largely as interchangeable “housekeeping” factors. However, growing evidence argues the opposite: many glycolytic enzymes exist as multiple isoforms with small sequence differences that yield distinct regulatory properties, kinetic behaviors, or subcellular localizations. This has been best described for mammalian hexokinase and pyruvate kinase isoforms, where tissue-specific expression determines whether glucose is routed toward catabolism or biosynthesis, and where distinct splice forms localize to mitochondria or cytosol to tune metabolic flux^21,22,58,59,64^. Despite this precedent, the field has lacked systematic *in vivo* studies that test whether different enzyme isoforms support specific physiological outputs underpinned by tissue-specific metabolism. GPI has been almost universally treated as a single, freely diffusing cytosolic enzyme, with little attention paid to isoform expression, tissue specificity, or subcellular targeting. Our work suggests that isoform variation can tune tissue-specific metabolic demand even for GPI-1, a highly conserved enzyme whose reversible reaction is classically viewed as operating near equilibrium with its substrates. These findings support a broader emerging view: that “housekeeping” enzymes harbor previously underappreciated regulatory axes capable of tuning carbon metabolism in a tissue-specific manner, and that resolving their isoform-level roles and localizations will likely reveal new layers of metabolic control across animals.

A key implication of our findings is that subcellular localization represents an underappreciated regulatory axis for directing glucose metabolism. Unlike irreversible committed steps, GPI catalyzes a near-equilibrium reaction in which net flux depends not on catalytic directionality, but on local metabolite availability. Thus, the spatial compartmentalization of GPI-1B on endomembrane structures provides a mechanism for tissue-specific metabolic control: by positioning GPI adjacent to distinct metabolic microenvironments, cells could potentially bias whether G6P is routed toward glycolysis or diverted into anabolic pathways such as the PPP. This concept is consistent with studies showing the PPP reactions are not homogeneously cytosolic, as ER-resident hexose-phosphate transporters and dehydrogenases initiate NADPH-generating reactions^65–70^. By localizing GPI-1B to these same structures, the germline could transiently enrich G6P at the ER to support PPP activity, biosynthesis, redox balance, and embryonic development. Conversely, somatic tissues lacking GPI-1B localization operate with a diffuse, cytosolic GPI-1A, better configured to support rapid glycolytic flux. Thus, our results highlight that for enzymes catalyzing reversible reactions such as GPI, subcellular localization could be a physiologically meaningful layer of metabolic regulation.

These findings also suggest mechanistic models by which organelle-associated pools of GPI-1B could shape metabolic flux. One possibility is that ER targeting localizes GPI upstream of ER-resident glucose-6-phosphate dehydrogenase and hexose-phosphate transporters, creating a spatially privileged route for G6P entry into the PPP. Another is that GPI-1B on the ER or ER-derived lipid droplets may reduce the compartmental availability of G6P for cytosolic glycolytic enzymes, restricting competition with phosphofructokinase and biasing flux toward anabolic metabolism. This model is consistent with growing evidence that metabolic enzymes form spatially organized assemblies capable of channeling intermediates, protecting unstable metabolites, or altering apparent reaction equilibria^52,55,71–74^. The observation that specific loss of GPI-1B expression impairs reproductive fitness, despite intact enzymatic function of GPI-1A, raises the possibility that organelle targeting of reversible enzymes may be a general strategy to tune pathway choice without altering transcription, translation, or protein abundance. These hypotheses are now experimentally tractable and invite future work to identify interacting partners of ER-localized GPI-1B, quantify compartment-specific sugar phosphate pools, and test whether organelle targeting of reversible enzymes sculpts metabolic wiring during development, stress, and disease.

## ACKNOWLEDGEMENTS

We thank the labs of Shaul Yogev, Marc Hammarlund and Pietro De Camili for sharing reagents and constructs. We thank Frank Schroeder, Bennett Fox and Camila Castellanos for their advice, resources and metabolomics analysis. We thank Dora Tang, Martine Ruer and Anjali Anilkumar for their efforts towards *in vitro* reconstitution of GPI-1 isoforms. We thank Diego de Mendoza, Andrés Binolfi and Carla Delprato for their lipidomics analysis into GPI-1 single isoform mutants. We thank Meng Wang and Dinghuan Deng for their time and resources performing stimulated Raman scattering analysis on GPI-1 single isoform mutants, and for the insight into their tissue-specific single cell RNA sequencing data. We thank Valerie Reinke for advice and insight into reproductive and germline defects we observed in *gpi-1* mutants. We thank Tim Schedl, Josh Bembenek and Barth Grant for their insights into subcellular GPI-1B localization in the germline. We also thank members of the Colón-Ramos Lab and Dick Goodman for feedback on figures and the manuscript. We also thank Stacy Wilson for technical support and training using the Airyscan imaging set-up as part of the Yale Neuroscience Imaging Core Facility. Some strains were provided by the CGC, which is funded by NIH Office of Research Infrastructure Programs (P40 OD010440). Some figures were created in BioRender. Metabolomics study and analysis (performed by QS, SS, RK) were done via a collaboration with the Yale Chemical Metabolism core facility. This work was supported by National Institutes of Health grants to DC-R (R35NS132156 and R01NS076558), A.J.M.W. (R35GM122502 and R01DK068429) and AW (K99AG083129). This work was also supported by the NSF GRFP to IG. MH was also supported by HHMI.

## METHODS

### *C. elegans* husbandry and genetics

All worm strains were grown on standard laboratory nematode growth media plates with *E. coli* strain OP50 as the sole food source and plates were kept at 20 °C. Unless explicitly stated, experiments were done with day 1 adults. The *C. elegans* Bristol N2 strain was used as the wild type control for all experiments. Recovered worms (from thawing or starved plates) were kept in well-fed conditions for a minimum of 3 generations before being used for experimentation. Worms with lethal/sterile mutations were maintained as heterozygotes with chromosomal balancers. The strain list at the end of the methods section describes these strains in more detail. Genetic rescue experiments were done with plasmids with the pttx-3g promoter, cDNA of the relevant sequence, and the *unc-54* 3’UTR sequence.

### Molecular Biology

#### Cloning and Injections

Gibson Assembly protocols were used to construct plasmids for microinjection and creation of transgenic *C. elegans* strains. Standard microinjection protocols were used to generate transgenic strains^75^, involving microinjection of a construct mix where the final concentration of all plasmid elements was no more than 100ng/μl.

#### CRISPR-Cas9 genome editing

CRISPR genome editing protocols were used to generate mutants and were based on previously established protocols^76^. Guide RNA sequences were designed by inputting genomic sequences near the site of the desired genome lesion or modification into the CRISPOR web tool (https://crispor.gi.ucsc.edu/crispor.py)^77^ and the Design custom gRNA web tool from IDT (<u>idtdna.com/site/order/designtool/index/CRISPR_SEQUENCE</u>) (Table 2). Designed guides were chosen based on similarity in evaluated efficiency and specificity between both tools. We evaluated efficiency and specificity using CRISPOR’s *Predicted Efficiency* and *Specificity Score* metrics, as well as IDT’s *On-target Potential* and *Off-target Risk* metrics. Guides were then ordered from Horizon Discovery (Dharmacon) as Edit-R Modified Synthetic crRNAs. A gRNA sequence to create the *dpy-10 (cn64)* allele was used as a co-CRISPR target to track possible successful modifications via Cas9. Genomic edits targeting single amino acids (*GPI-1B-expressing animals/ GPI-1B only* mutants) utilized one guide RNA, while larger deletion mutants (*GPI-1A-expressing animals/ GPI-1A only* mutants and *gpi-1* KO mutants) required two guide RNAs, one 5’ of the deletion and one 3’ of the deletion. Corresponding homology templates were designed in SnapGene such that there was a 35 base pair overlap on both sides of the Cas9 cut site. Designed templates were then ordered from the Keck Oligonucleotide Synthesis facility at the Yale School of Medicine as linear, single stranded oligonucleotides (Table 2). To make the CRISPR injection mix, purified Cas9 protein, tracrRNA, *dpy-10* gRNA, and the custom gRNA of choice were mixed on ice in the volumes and concentrations below in order, mixed gently with pipetting, and incubated at 37 °C in a thermocycler for 10 minutes. Note: some alleles, such as point mutations, only require one gRNA, while larger deletions require two flanking guides.

**Table 1:**
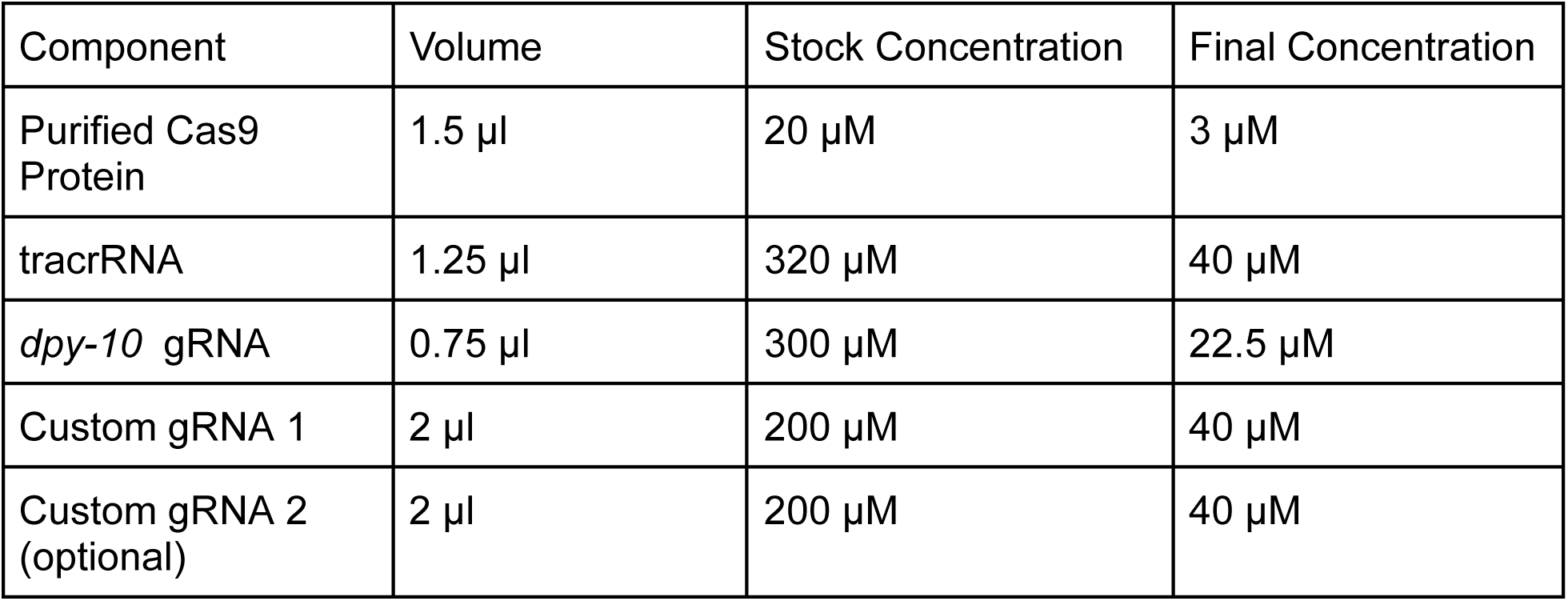
Formulation of CRISPR-Cas9 injection mix for designing custom alleles.

**Table 2:**
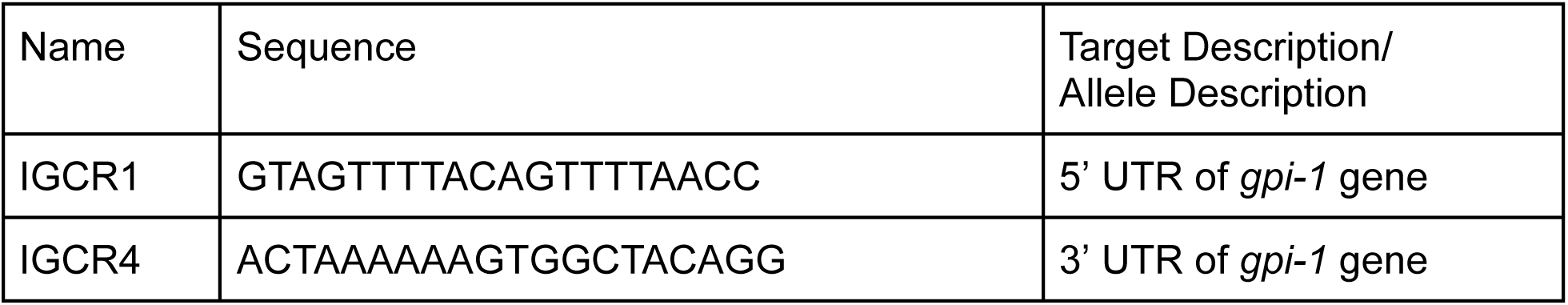
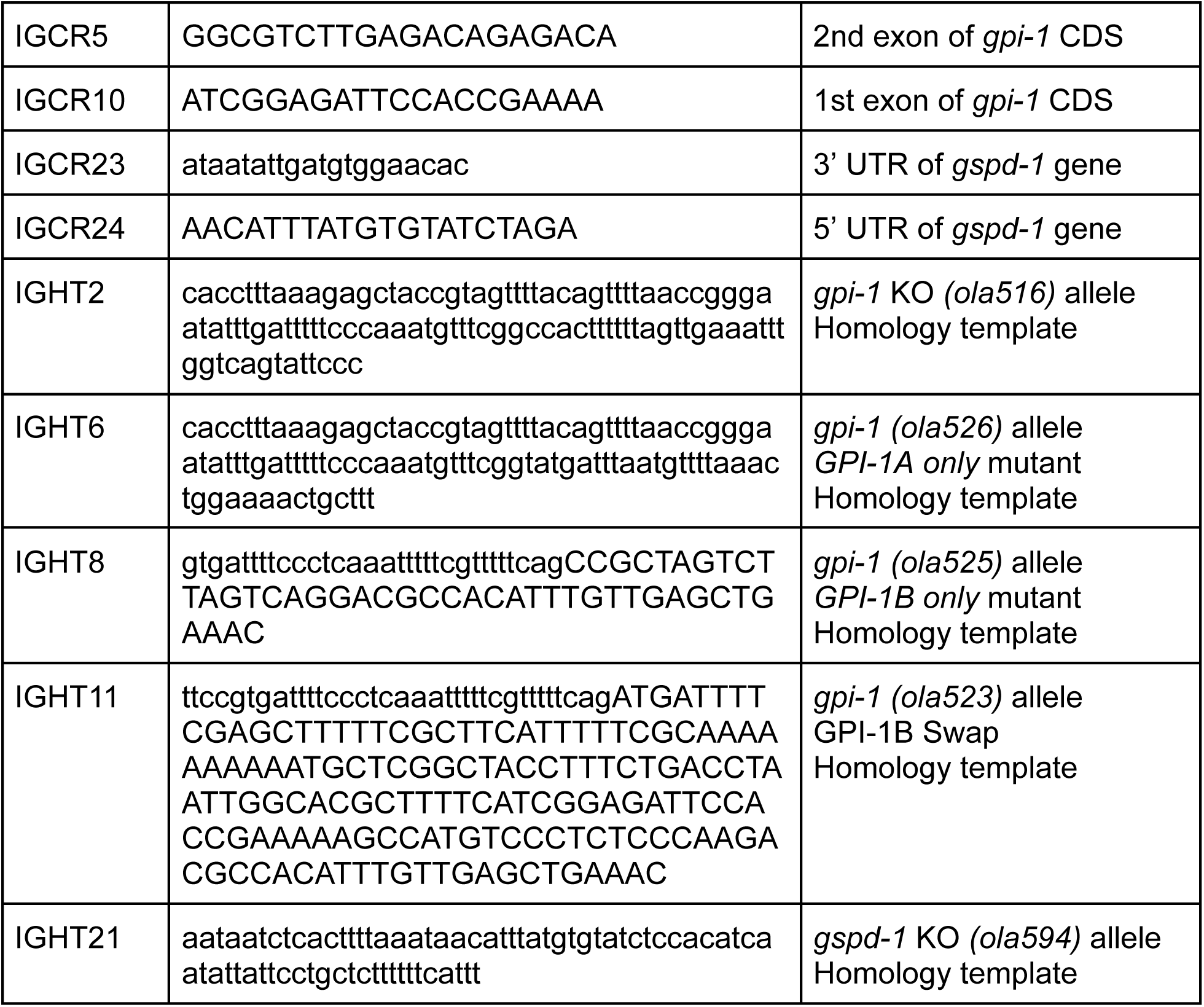
List of CRISPR gRNAs (IGCRs) and homology templates (IGHTs) sequences used in relevant experiments with description of target or corresponding allele for each.

Following incubation, the mix of Cas9 and guide RNAs can be used at room temperature, although working in an ice bucket is preferable. Next, 1μl of single stranded homology template for the desired edit and 0.5 μl of the *dpy-10 (cn64)* allele homology template were each added at a concentration of 10 ng/μl, for a final concentration of 1 ng/μl and 0.5 ng/μl respectively. The total volume of the mix was then made to 10μl via the addition of nuclease-free water. Finally, the mixture was gently mixed by pipetting, and spun in a microcentrifuge at 12,000 x g for 10 minutes to pellet any dust or debris - an essential step for successful microinjection. The top 5μl of the solution was then loaded into a capillary-based microinjection needle and injected into the gonads of day 1 adult animals. Following injection, F1 progeny were screened for the roller phenotype (heterozygous phenotype for the *dpy-10 (cn64)* allele) and then separated onto individual plates for subsequent genotyping for the allele of interest. Upon confirmation of the allele in individually plated worms from the F1 generation, select worms were outcrossed to eliminate the *dpy-10* allele and self-crossed to make the desired allele homozygous.

CRISPR-Cas9 was similarly used by SunyBiotech to generate the *gpi-1 (ola528)* allele, which features a full-length GFP inserted at the 3’ end of the *gpi-1* CDS. SunyBiotech also made the DCR29 (G001-Pitr-1-SL2-gpi-1cDNA-GFP-unc-54 3’UTR-EG6699 or PHX2188) DA9 Single copy insertion of GPI-1B::GFP strain via MOSCI.

### Cell culture and Transfections

hTERT-RPE-1 (RPE-1) cells were grown in DMEM/F12 medium (Thermo Fisher Scientific) supplemented with 10% FBS, 100 U/mL penicillin, 100mg/mL streptomycin and maintained at 37°C in humidified atmosphere at 5% CO_2_. 2mM glutamax (Thermo Fisher Scientific) was added to all media for RPE-1 cells. Cell lines were routinely tested and always resulted free from mycoplasma contamination. Transient transfections were carried out on cells that were seeded at least 8 hours prior. All transfections of plasmids used FuGENEHD (Promega) to manufacturers specifications for 16-24 hours in complete media without antibiotics.

### Mounting and Microfluidics

Worms were mounted for imaging on 10% agarose pads dissolved in water. Worms were placed in the center of 2.5 μl of M9 mixed 1:1 with polystyrene 0.1-micron beads (Polybead® Microspheres - Cat# 00876-15) or in 2.5 μl of 10 mM levamisole dissolved in M9 that was pipetted onto agarose pads. A 22x22mm No. 1.5 glass coverslip was then gently placed on the agarose pad. For induction of acute hypoxia, modified PDMS microfluidics devices enabling precision gas flow of nitrogen (hypoxia) and breathing air (normoxia) were used as described previously^54,55^. Agarose pads were affixed to the top of the PDMS devices where the gas exchange occurred, which when topped with a glass coverslip allowed for acute and local induction of hypoxic stimuli.

### Microscopy and analysis

Microscopy experiments featuring live *C. elegans* were conducted almost exclusively using a Nikon Ti2 + CSU-W1 spinning disk confocal microscope. Images were captured using a Hamamatsu Ocra-Fusion BT CMOS camera at 16-bit pixel depth. Sample excitation was done according to a given strain’s expression of fluorescent reporter, either at 405, 488 or 561 nm wavelength via 50 mW lasers for each. Ratiometric imaging of the HYlight FBP biosensor was done via alternating excitation by the 488 and 405 nm lasers without changing the 525 nm emission filter. Laser settings for AIY HYlight imaging done at 10x magnification were set to 8% laser power for the 488 nm laser and 2% laser power for the 405 nm laser. Pan-neuronal HYlight imaging was performed with 8% laser power for the 488 nm laser and 4% laser power for the 405 nm laser. Both imaging conditions were done with each laser set at the same exposure time which ranged from 10 to 200 ms. Hypoxic induction during experiments was done via manually switching between gas tanks at the indicated time on relevant data graphs and charts. Normoxic and hypoxic (-O_2_) comparisons of AIY HYlight reflected values representing the ratio at the beginning of the normoxic stimulus and the end of the hypoxic stimulus. Fiji analysis software was used for analysis and for the construction of maximum intensity projections^78^. For representative images of FBP HYlight ratio, the 488-nm excitation channel was divided by the 405-nm channel to create a ratiometric image. The LUT was set to mpl-magma and the ratio bounds were scaled to the ranges depicted in figure images. Quantification of HYlight images was done similarly to previous protocols^15,41^. For AIY HYlight images, background subtraction was applied to both the 488- and 405-nm channel and a region of interest was drawn around the cell soma for each channel. The mean pixel value of this ROI was then used as the value for each channel, and the ROI ratio for each cell was calculated by dividing the 488-nm ROI value by the 405-nm ROI value. The FiLa lactate biosensor was codon-optimized to C. elegans, an intron was inserted, and the FiLa sequence was fused to Mscarlet-I3 via a T2A sequence. The sensor was excited by the 405-laser using a 525 nm emission filter. The Mscarleti3 fluorophore was excited at 561nm using a 605 nm emission filter. Laser settings for FiLa imaging done at 10x magnification were set to 20% laser power for the 405 nm laser and 1% laser power for the 561 nm laser, with an exposure time of 120ms. Ratiometric quantification of FiLa was done on the Fiji analysis software, similarly to previous protocols (cite HYlight paper) using 1 divided by 405nm/561nm. One-way ANOVA was used for statistical analysis.

Electron microscopy was used to visualize the localization of GPI-1B pan-neuronally expressed in the *C. elegans* nervous system. High-pressure freezing, freeze substitution, and sectioning were all performed as previously described^79–82^. For immuno-EM, sections of 50 nm were collected on nickel slot grids covered with Formvar (EMS). Grids were incubated at 20°C on 50 μl droplets of 0.05 M glycine PBS for 5 min, 1% BSA and 1% CWFS gelatin in PBS for 20 min, anti-GFP rabbit polyclonal (1:20 10ug/ml in 0.3% BSA and 0.3% CWFS gelatin in PBS, ab6556 Abcam) overnight at 4°C and then 60 min at 20°C, 6 PBS washes over 30 min, Protein A Gold conjugated to 10 nm gold (1:75 in 0.3% BSA and 0.3% CWFS gelatin in PBS, University Medical Center Utrecht) for 60 min, 6 PBS washes over 30 min, 2% glutaraldehyde in PBS for 5 min, 3 water washes for 10 s. After drying, grids were post-stained in 2% uranyl acetate for 4 min, and lead citrate for 1 min. Images were acquired on TALOS L120 (Thermo Fisher) equipped with a Ceta 4k × 4k CMOS camera.

For Airyscan imaging, worms were mounted on a Zeiss LSM880 microscope equipped with an Airyscan detector and imaged with a 63X NA 1.4 oil objective.

### Live Cell Imaging and Immunofluorescence

For all live cell microscopy cells were seeded on glass-bottom mat-tek dishes (MATtek corporation) 5500/cm^2^ in complete media. Transfections were carried out as described above. Spinning-disk confocal imaging was performed 16-24 h post transfection using an Andor Dragonfly 200 (Oxford Instruments) inverted microscope equipped with a Zyla cMOS 5.5 camera and controlled by Fusion (Oxford Instruments) software. Laser lines used: DAPI, 440nm; GFP, 488; RFP, 561; Cy5, 647. Images were acquired with a PlanApo objective (60x 1.45-NA). During imaging, cells were maintained in Live Cell Imaging buffer (Life Technologies) in a cage incubator (Okolab) with a humidified atmosphere at 37°C.

### Developmental and reproductive assays

#### Age synchronization via egg lay

For experiments comparing developmental dynamics, body length and onset of egg laying, worms were synchronized via a 1-4 hour egg lay. Briefly, 10-20 L4s of each strain to be tested were isolated onto a standard NGM plate seeded with OP50.

Approximately 16 hours later, the now day 1 adult animals were placed onto a separate seeded NGM plate and allowed to lay eggs for 1-4 hours. Following the egg lay, worms were removed and returned to their original plate, at which point, the newly laid embryos would be considered to be “0 hours old” and monitored for subsequent experimentation.

#### Quantification of body length and developmental milestones

To measure body length over time, newly laid embryos from age-synchronized strains were monitored and imaged starting 24 hours after the end of the egg lay. Plates were then imaged in 8-16 hour increments using the WormLab imaging system equipped with a Basler acA2440- 35mm monochromatic sensor with an infrared filter as previously described^83^. A 10 mm mark was created on each worm plate and used to calibrate length on captured images for subsequent measurement of worm body length over larval development.

To measure when worms reached developmental milestones such as entering the L4 stage and the onset of egg production, age-synchronized strains were scored on a benchtop stereomicroscope every 8 hours starting 48 hours after being laid. L4 stages were determined as described previously and were noted across different strains and genotypes^84^. Onset of egg production was determined by mounting synchronized worms for 60x brightfield imaging on the aforementioned spinning disk confocal microscope every 8 hours starting 48 hours after being laid. In a given sample and timepoint, worms were scored for presence of *in utero* embryos, marking the onset of egg production.

This proportion of animals that had started egg production at a given timepoint to all animals scored was then calculated and represented over time following the initial egg lay.

#### Brood size analysis

Brood size analysis was done as previously described^85^. Briefly, to measure the brood size of animals across different genetic backgrounds, 10 L4 worms were transferred onto individual plates for each genotype and labeled as Day 1; Worm1; Genotype 1, Day 1; Worm X; Genotype X, etc. 24 hours following the first transfer, each worm was transferred to a plate marking the same worm and genotype, but with Day 2 as the label. This was done until Day 5, being sure to only transfer the specific worm from the original Day 1 plate and not any unhatched embryos or progeny. This resulted in five plates for each worm in a given genotype. Unhatched embryos and live progeny on each plate were counted 48 hours after each transfer until all five plates for each worm and genotype were recorded. The sum total of a worm’s live progeny was then calculated for each worm and averaged across plates from the same genotype in order to compare the effect of each genotype on brood size or hatched embryos. One-way ANOVA was used for statistical analysis. We occasionally observed that *pfk-1.1* KO animals would demonstrate an embryonic lethality phenotype, laying similar quantities of non-hatching eggs as hatched eggs measured in this study. We found that *pfk-1.1* KO animals could be maintained with the described metrics of reproductive fitness and accelerated development if grown on newly made NGM plates with thicker OP50 bacteria lawns. We hypothesized that this may represent a specific dietary regimen that enables *pfk-1.1* KO animals to metabolically adapt, as well as a sensitivity towards high salt concentrations associated with older NGM plates that have undergone water evaporation. We note that when these mutants are grown in conditions that vary

### Swimming Assay

Swimming behavior analysis was performed by imaging with the WormLab imaging system equipped with a Basler acA2440- 35mm monochromatic sensor with an infrared filter and a computer running WormLab 2023.1.1 software (MBF Bioscience LLC, Williston, VT USA). As described previously^52^, worms were placed in a 20 μl drop of M9 physiological buffer (vehicle) or 20 μl of 1 mM sodium azide (NaN_3_) to disrupt mitochondrial oxidative phosphorylation. Worms were incubated for 5 minutes and were then imaged for 30 seconds to 1 minute at 15 fps to measure swimming behavior in each buffer condition. The WormLab analysis software was then used to build tracks of each worm’s movement over the duration of the video. Manual mending of tracks was occasionally performed when appropriate when the tracking software registered the same animals’ movement as more than one individual track. Swimming metrics were then automatically calculated for each track using the WormLab analysis platform based on the CeleST computer vision software^50,51^. The wave initiation rate, referring to the number of body waves initiated from either the head or the tail per minute, was then averaged across a given genotype and compared to wild type controls for each buffer condition. One-way ANOVA was used for statistical analysis.

### Targeted/untargeted metabolomics platform

#### Sample preparation

Approximately 10,000 *C. elegans* adult hermaphrodites per biological replicate were grown on 3-4 10 cm plates seeded with OP50. Once worms reached the first day of adulthood, worms were washed off multiple times with M9 physiological buffer, resuspended in ice cold quench buffer (150 µL/well, composed of 20% methanol, 3 mM sodium fluoride, 32 µM D_8_-phenylalanine as internal standard and 0.1% formic acid) and snap frozen in liquid nitrogen. Worm pellets were then lyophilized in Eppendorf tubes. Samples were prepared by resuspending the cell pellet in 10% acetonitrile solution with D_4_-taurine (25 µM) as a second internal standard. 5 µL of the supernatant was injected for each analysis mode on the mass spectrometer.

#### Chromatography

Two columns were used separately: a Thermo Scientific Hypercarb™ Porous Graphitic Carbon HPLC Column (100 x 4.6 mm, 3 µm) and a Phenomenex Kinetex F5 Core-shell HPLC column (100 x 2.1 mm, 2.6 µm).

Hypercarb column: Separations were performed with 1 mL/min linear gradients as indicated (Table 1). Mobile phase A: 15 mM ammonium formate, 0.03% acetyl acetone and 0.1% formic acid; mobile phase B: 60% ACN, 35% IPA, 15 mM ammonium formate and 0.1% formic acid. Column temperature was 50 °C and autosampler temperature was 5 °C.

**Table 1:**
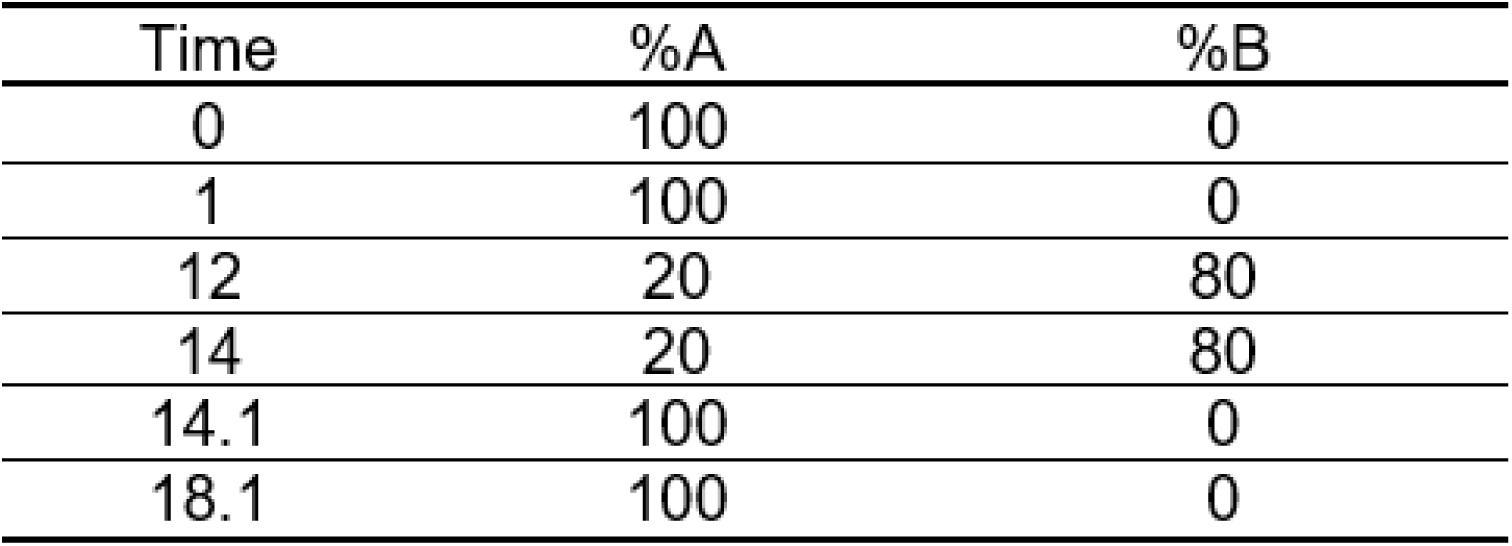
Hypercarb column chromatographic separation.

Kinetex F5 column: Separations were performed with 0.3 mL/min linear gradients as outlined (Table 2). Mobile phase A: 95% water, 5% acetonitrile and 0.1% formic acid; mobile phase B: 95% acetonitrile, 5% water and 0.1% formic acid. Column temperature was 30 °C and autosampler temperature was 5 °C.

**Table 2:**
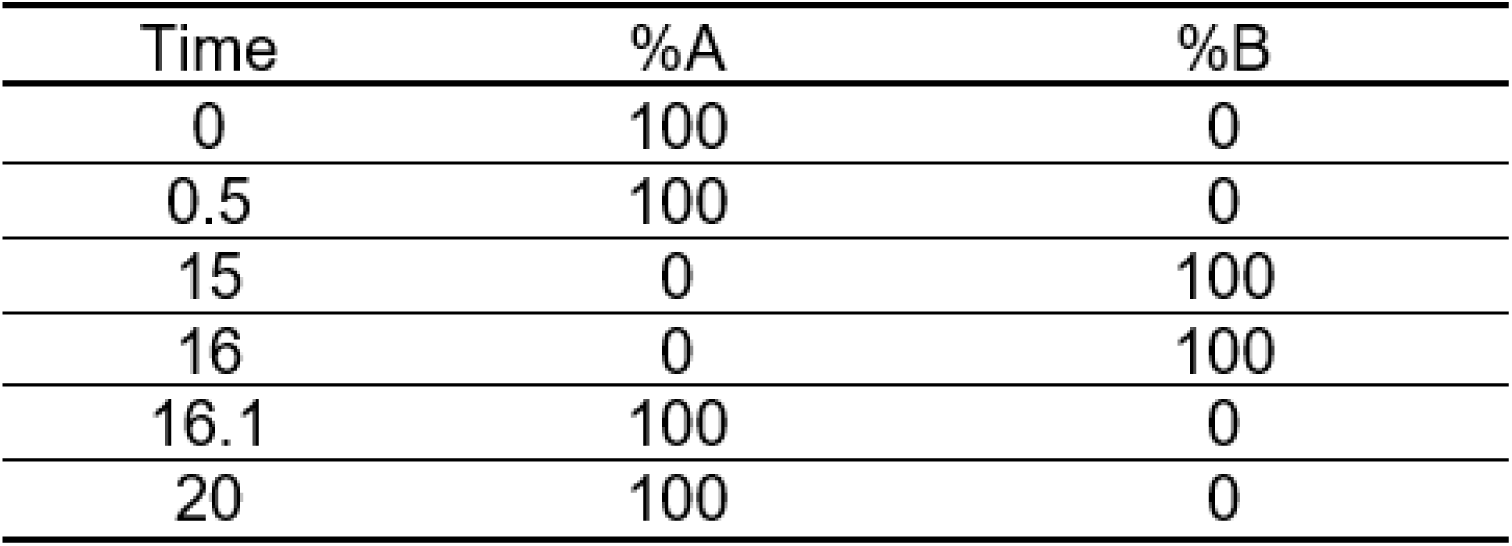
Kinetex F5 column chromatographic separation.

#### Mass spectrometry

Data were analyzed using the Sciex TripleTOF 6600 collected using an information-dependent analysis (IDA) workflow consisting of a TOF MS scan (200 msec) and a high-resolution IDA experiment (70 msec each) monitoring 10 candidate ions per cycle. Former target ions were excluded after 2 occurrences for 5 sec and dynamic background subtraction was employed. The mass range for both TOF MS and IDA MS/MS scans was 60-1000 with RP column and 70-1000 with Hypercarb column.

The ion source conditions were as follows; Ion spray voltage = 5000 V for positive mode and -4500 for negative mode, ion source gas 1 (GS1) = 50, ion source gas 2 (GS2) = 50, curtain gas (CUR) = 30, temperature (TEM) = 400 °C with F5 column and 500 °C with Hypercarb column. Compound dependent parameters for all four modes were: declustering potential (DP) = 35, collision energy (CE) = 30, collision energy spread (CES) = 20.

#### Data processing

El-MAVEN software (Elucidata.io) was used for peak picking and curation from house built targeted and untargeted libraries. Targeted libraries used the commercial standard kit of ∼ 600 metabolites (IROA Technologies) in all 4 modes of analyses (2 LC systems, positive & negative ion acquisition modes) yielding reference data (molecular ions and retention times) with wide coverage of the endogenous metabolome. Untargeted libraries contain ∼2,700 metabolites drawn from the KEGG database, and the top 5 candidates with the highest intensity throughout a sample run for each metabolite were automatically curated.

Data were normalized by the sum of all targeted metabolites within each sample in each mode and then log transformed. Sample quality was assessed using two internal standards, D_8_-phenylalanine and D_4_-taurine to identify any outliers to be excluded from the downstream analysis. Next, the normalized data from all four modes were merged. If a metabolite appeared in more than one mode, the one with the best intensity was chosen.

PCA, PLS-DA and heat map on the merged targeted data were performed using MetaboAnalyst 5.0 (https://www.metaboanalyst.ca/). Differential expression analysis was performed using the metabolomics application in Polly (polly.elucidata.io). Normal p values and log2FC data was uploaded in Shiny GATOM (https://artyomovlab.wustl.edu/shiny/gatom/) to generate an optimized metabolic network displaying ∼60 of the most changing and closely connected metabolites. The up-regulated and down-regulated metabolites in the study group were then used separately for pathway enrichment analysis in MetaboAnalyst 5.0 (https://www.metaboanalyst.ca/).

#### Conservation of metabolic genes calculation

Multiple sequence alignments were generated for *gspd-1*, *gpi-1*, *pfk-1.1* and *pfk-1.2* by comparing with annotated orthologs from the species *Drosophila melanogaster*, *Danio rerio, Rattus norvegicus, Mus musculus,* and *Homo sapiens*. Alignments were generated using the MUSCLE algorithm^86^, and a weighted consensus conservation score for each gene was calculated using Shannon entropy^87^. This alignment was then visualized on the AlphaFold prediction^88^ for each of the *C. elegans* orthologs by coloring by conservation at each residue using ChimeraX^89^, using a range of Z-scores from -2 to 2. A higher Z score reflects residues that are more highly conserved across species.

#### Metabolic Network Modeling

Single-cell RNA-seq data from wild-type Day 1 adult C. elegans^38,42^ were aggregated into pseudo-bulk expression profiles (TPM) for every cell type following established procedures and used for metabolic modeling. These data were integrated with the C. elegans genome-scale metabolic network model iCEL1314 using the MERGE framework^38^.

For each cell type, a feasible network-wide flux state was inferred using the iMAT++ algorithm. Gene expression values were first converted into discrete activity states using CatEx^38^. Simulations were performed using iCEL1314 in its standard form^38^ with modifications to relax the biomass objective (i.e., instead of requiring full biomass synthesis, sink reactions were introduced for major biomass precursors to allow partial biosynthetic activity). Standard exchange constraints were applied to permit uptake of metabolites that can be supplied systemically or are necessary to sustain flux. At the same time, biologically implausible transports were discouraged by assigning the exchange of intermediate metabolites (e.g., TCA cycle intermediates) and compounds unlikely to be available in vivo to the low-expression category, while metabolites not present in the endogenous C. elegans metabolome^90^ were excluded using hard constraints. Following optimization, flux variability analysis (FVA) was performed to determine the allowable flux range for each reaction.

To quantify reaction-level flux capacity, we applied the eFPA algorithm^39^ using the same constrained version of iCEL1314. Expression information was incorporated within a local network neighborhood defined by default distance parameters^39^. To limit the use of unlikely exchange fluxes, boundary reactions associated with discouraged metabolite uptake or secretion were penalized (penalty factor = 25). Finally, reaction bounds were adjusted based on FVA results to enforce directional feasibility, ensuring that eFPA-derived flux potentials remained within the solution space defined by iMAT++.

**Table 3:**
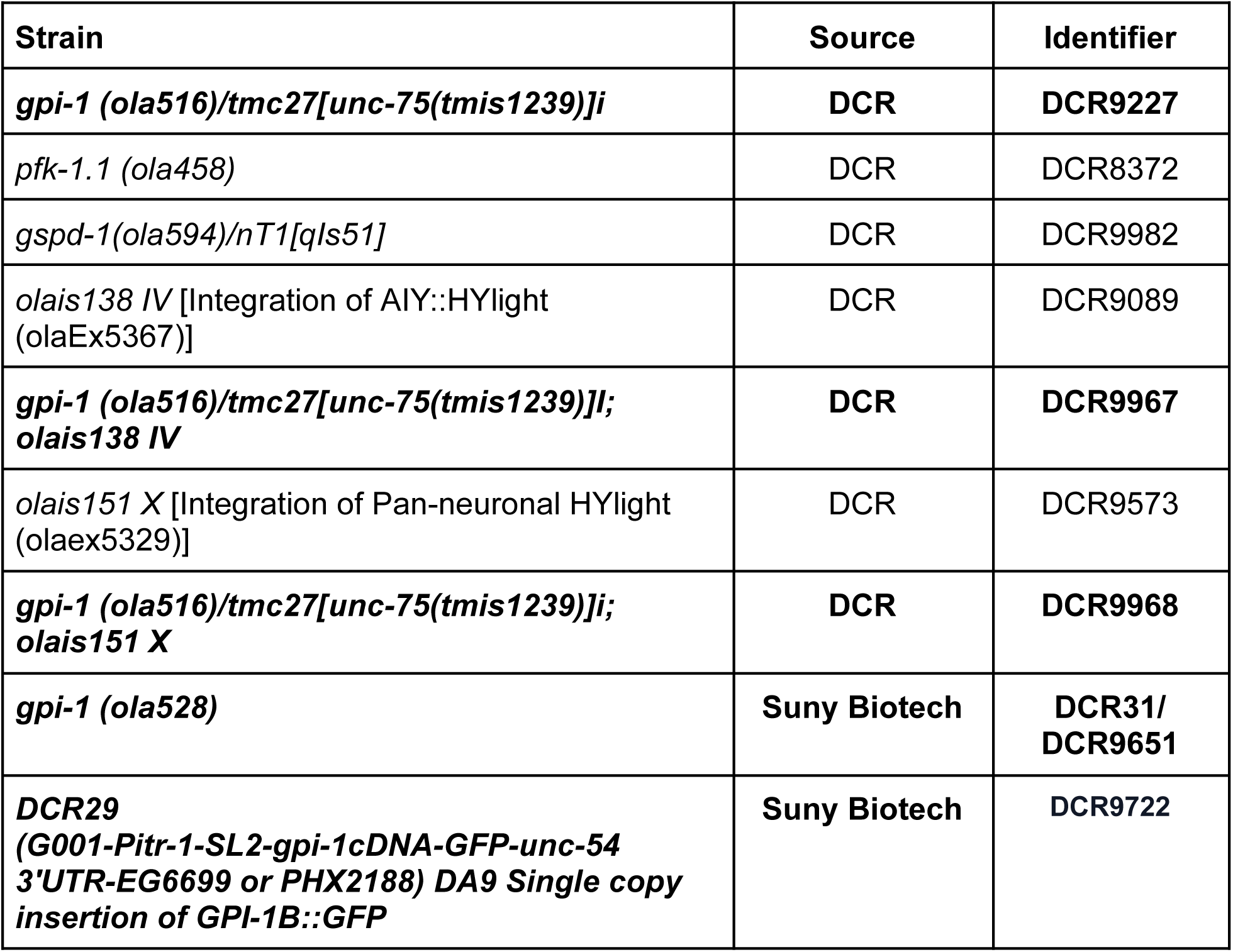
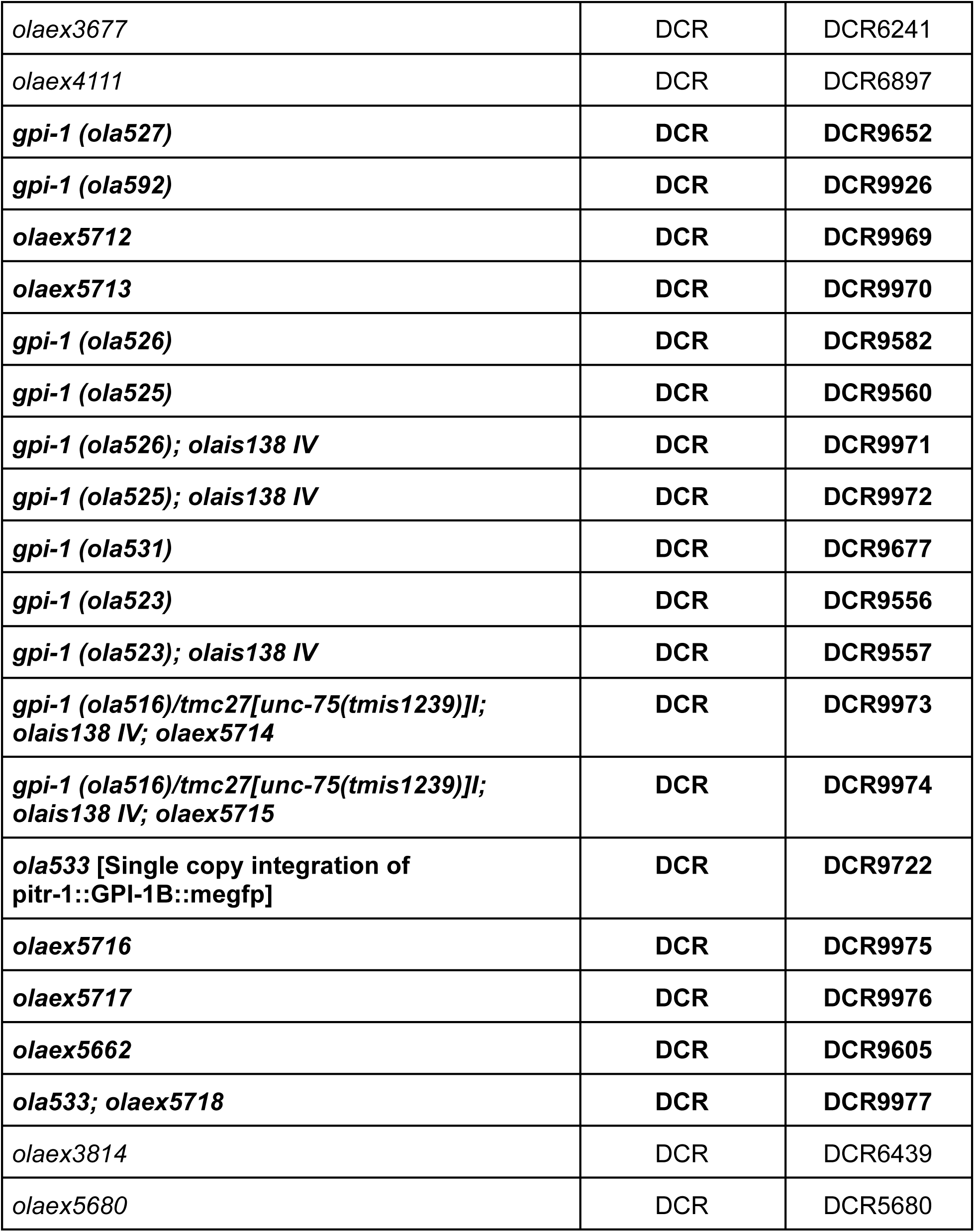
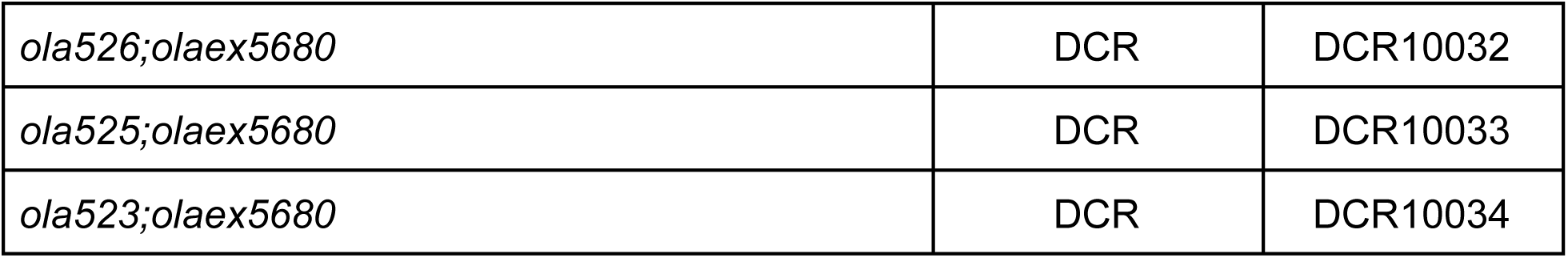
Nematode Strains. List of *C. elegans* strains used in this study. CGC is the Caenorhabditis Genetics Center, DCR is the Colón-Ramos Lab and NBRP is the National Bioresource Project (Shohei Mitani Lab strains). Strains specifically generated for this study are bolded.

---

**Supplemental Figure 1 (S1).**
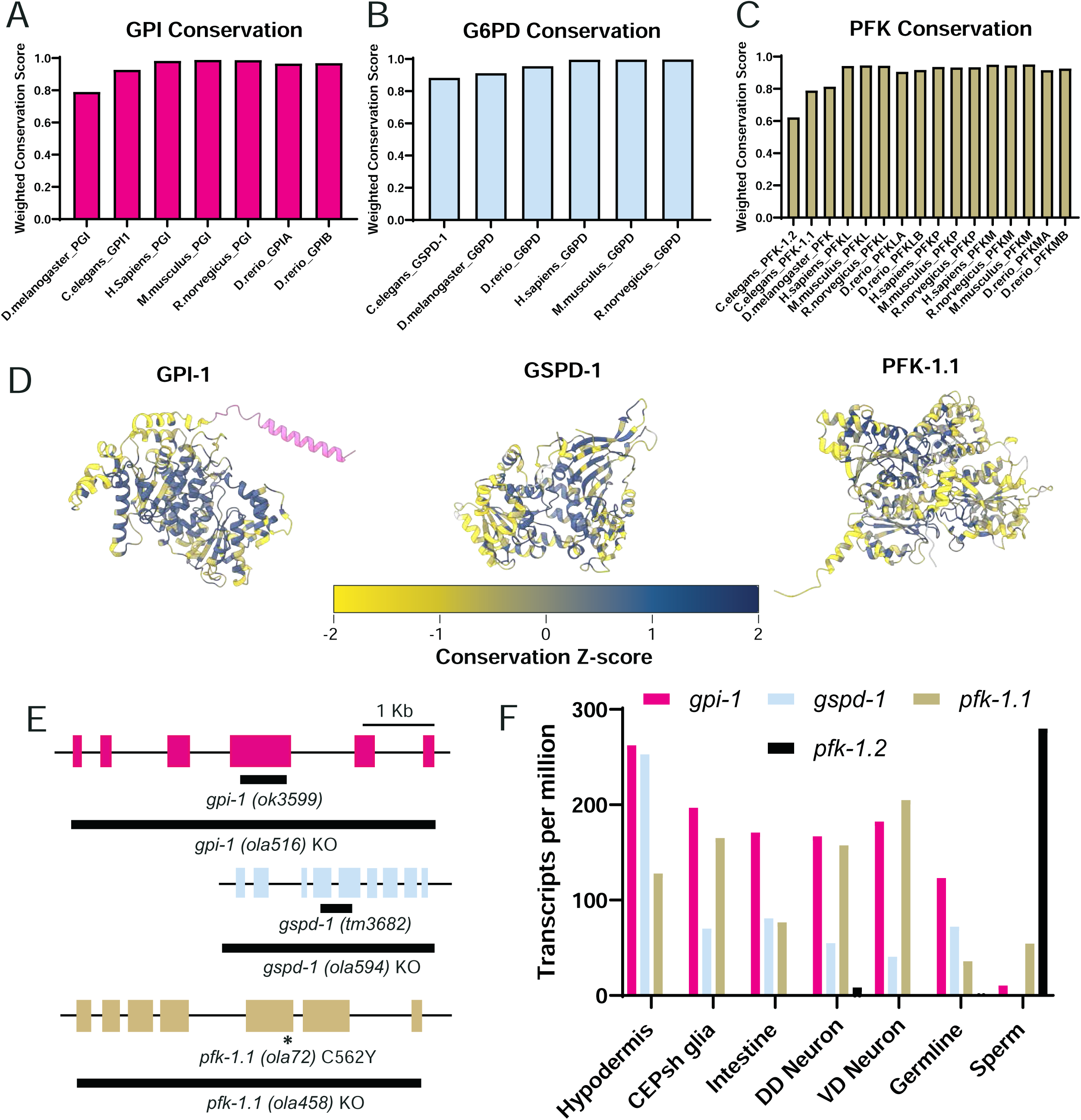
**(A-C)** A weighted conservation score was calculated relative to a consensus sequence generated by a multiple sequence alignment of known orthologs of GPI, G6PD, and PFK, respectively; the species *D. melanogaster, D. rerio, R. norvegicus, M. musculus, H. sapiens*, and *C. elegans* were compared. **(D)** AlphaFold predictions for the *C. elegans* proteins GPI-1, GSPD-1, and PFK-1.1 are colored by the conservation Z-score for each residue from each multiple sequence alignment. Color scale is shown below. In GPI-1, the unique GPI-1B exon at the N-terminus is colored in pink. **(E)** Schematics of the *C. elegans gpi-1, gspd-1* and *pfk-1.1* gene with correspondent allele names and CRISPR-deleted regions in respective gene knockouts denoted by black bar. **(F)** Expression level (in transcripts per million) of *gpi-1, gspd-1, pfk-1.1* and *pfk-1.2* genes across *C. elegans* tissues. *pfk-1.2* is an alternative variant of *pfk-1.1* that is expressed almost exclusively in sperm. TPM values from adult CeNGEN dataset^101^.

**Supplemental Figure 2 (S2):**
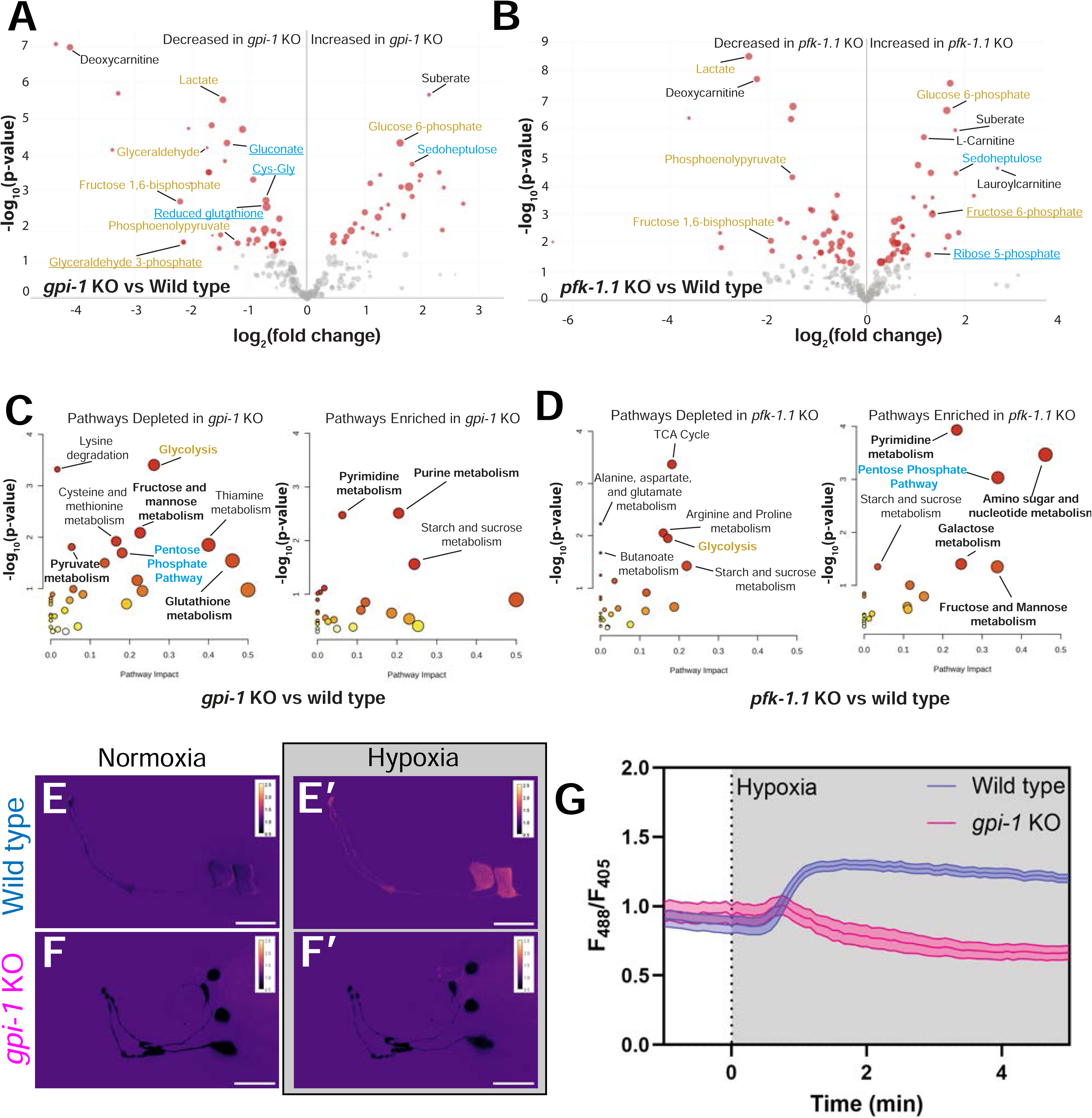
**(A - B)** Metabolomics analysis demonstrating differential metabolites between *gpi-1* KO (A) and *pfk-1.1* KO (B) relative to wild type controls. **(C-D)** Pathway analysis for *gpi-1* KO (C) and *pfk-1.1* KO (D) animals relative to wild type controls shows pathways depleted or enriched in each genetic background. **(E-F)** Ratiometric image of AIY HYlight in wild type (E) and *gpi-1* KO animals (F) and responses to acute hypoxic stimulus (E’ and F’). **(G)** Quantification of AIY HYlight response to acute hypoxia in wild type (blue) and *gpi-1* KO animals (pink).

**Supplemental Figure 3.**
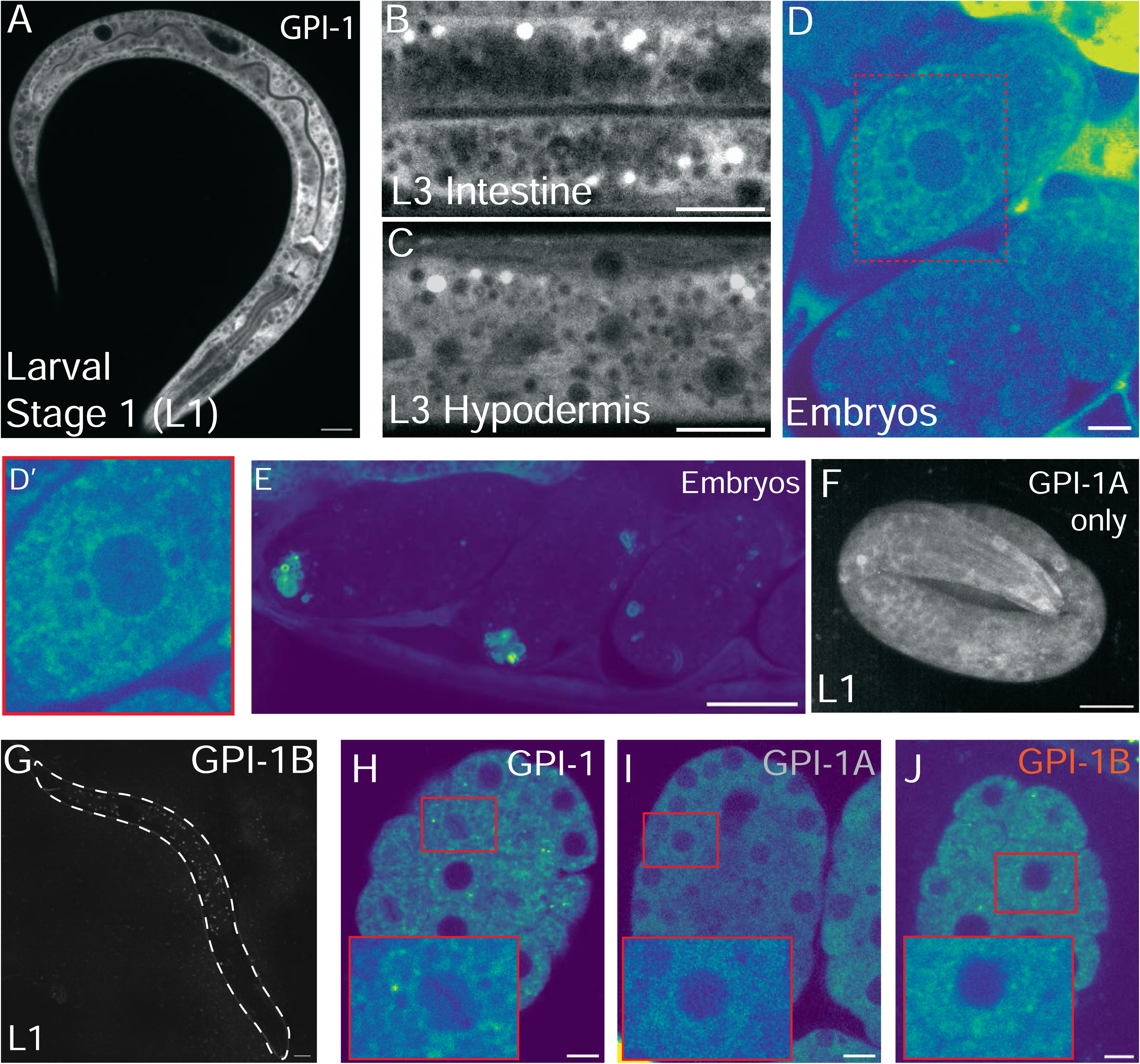
**(A)** GPI-1::GFP expression in larval stage 1 (L1). Scale bar - 10 microns. **(B-C)** Subcellular expression patterns of GPI-1 in intestine (B) and hypodermis (C) in L3 animals. Note that the bright spots represent autofluorescent gut granules and not GPI-1 localization. Scale bar - 10 microns. **(D)** GPI-1 expression in embryos with corresponding zoom-in image shows subcellular membranous perinuclear localization (zoomed-in in D’). Scale bar - 5 microns. **(E)** Subcellular localization of GPI-1 to membranous structures in embryos. Scale bar - 20 microns **(F)** *GPI-1A only* animals showing expression of GPI-1A early in larval development, demonstrating global expression across somatic tissues. Scale bar - 10 microns **(G)** *GPI-1B only* animals showing expression of GPI-1B early in larval development, demonstrating a lack of somatic tissue expression. Animal silhouette traced to facilitate visualization. Scale bar - 10 microns. **(H-J)** Expression of GPI-1 in laid embryos of wild type (H), *GPI-1A only* animals (I) and *GPI-1B only* animals (J) showing differential subcellular localization patterns. Zoom-in images in red. Scale bars - 5 microns.

**Supplemental Figure 4.**
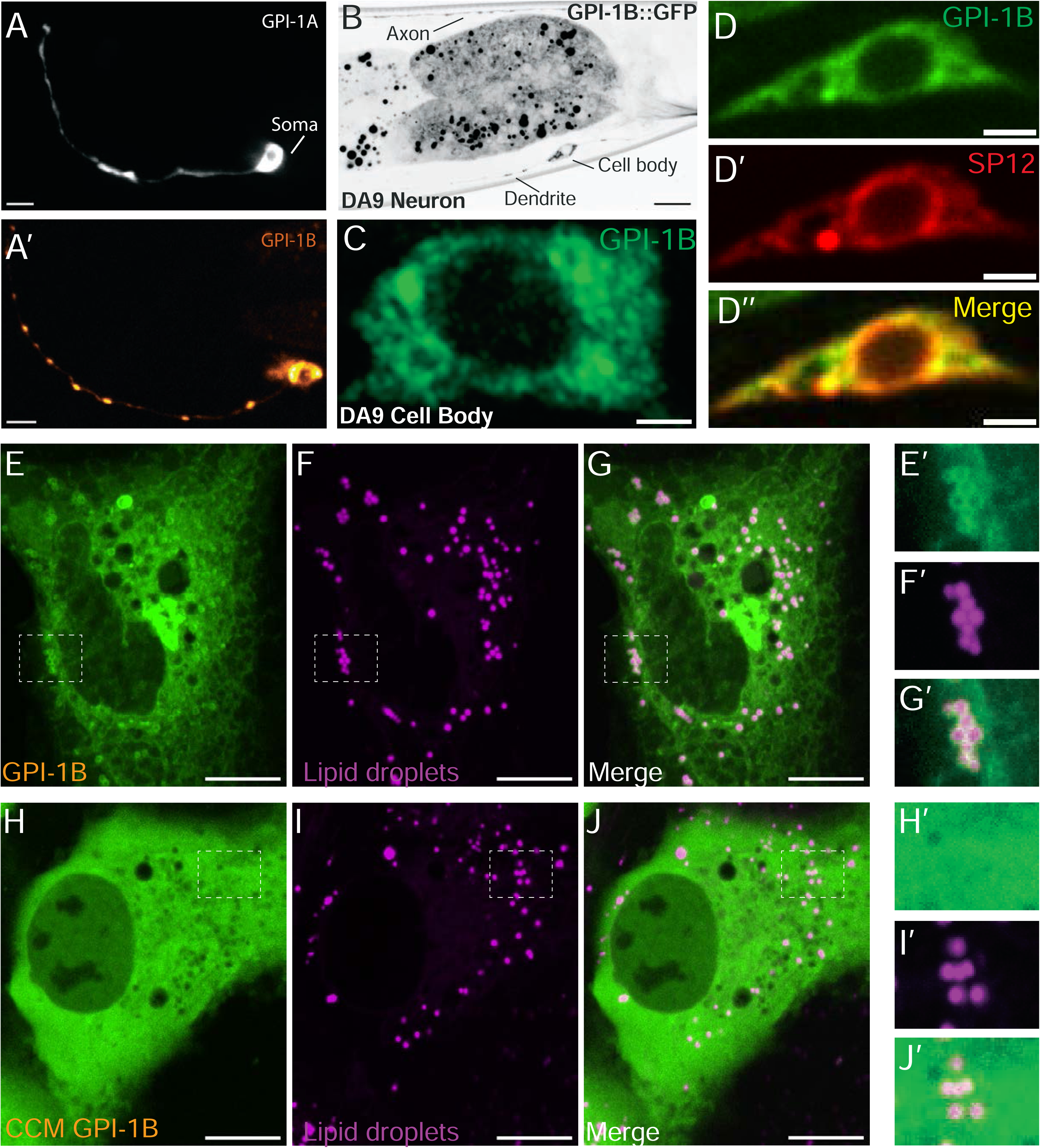
**(A)** Exogenous cDNA expression of GPI-1A (A) and GPI-1B (A’) in AIY neurons demonstrating localization pattern differences between GPI-1 isoforms. Scale bar - 5 microns. **(B)** Single copy expression of GPI-1B::GFP in the DA9 motor demonstrating localization in the neuron cell soma as well as the dendrite and axonal compartments. Scale bar - 10 microns **(C)** Superresolution Airyscan image of GPI-1B::GFP in the DA9 cell soma exhibiting localization to membranous and perinuclear structures. Scale bar - 1 micron. **(D)** Single copy expression of GPI-1B::GFP in the DA9 motor neuron cell soma with co-expression of general ER marker SP12 (D’) and merge image demonstrating colocalization (D’’). Scale bar - 5 microns. (**E)** Expression of GPI-1B in retinal pigmental epithelial (RPE-1) cells with inset image (E’). **(F)** Localization pattern of lipid droplets dyed by BODIPY 558/568 C12 stain with inset image (F’). **(G)** Co-localization of GPI-1B and lipid droplets in RPE-1 cells showing GPI-1B localization to outer lipid droplet membranes - inset image (G’). **(H)** Expression of CCM GPI-1B showing diffuse localization in RPE-1 cells, inset image (H’). **(I)** Lipid droplet marker in cells expressing CCM GPI-1B, inset image (I’) **(J)** Colocalization between CCM GPI-1B and lipid droplet marker showing lack of subcellular colocalization between markers in inset image (J’). (E - J) Scale bar - 10 microns.

**Table S1.**
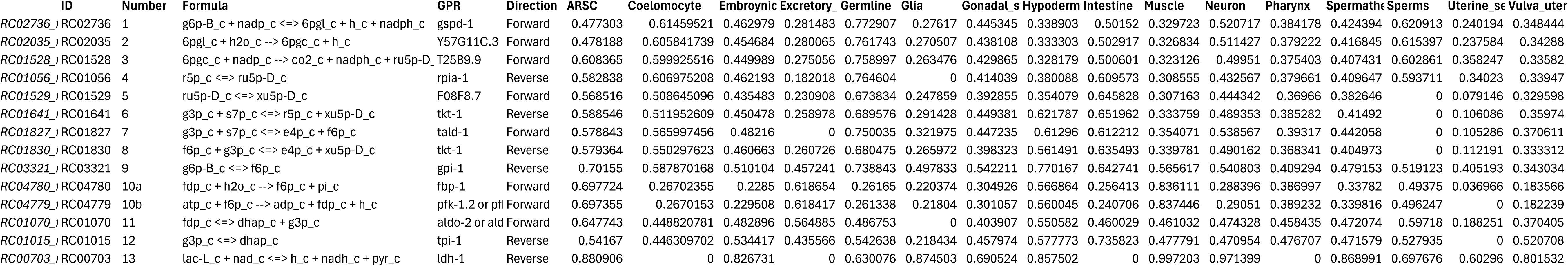

